# Active regulation of the epidermal growth factor receptor by the membrane bilayer

**DOI:** 10.1101/2025.08.14.670284

**Authors:** Shwetha Srinivasan, Xingcheng Lin, Xuyan Chen, Raju Regmi, Bin Zhang, Gabriela S. Schlau-Cohen

**Affiliations:** Department of Chemistry, Massachusetts Institute of Technology, Cambridge, Massachusetts, USA

## Abstract

Cell surface receptors transmit information across the plasma membrane to connect the extracellular environment to intracellular function. While the structures and interactions of the receptors have been long established as mediators of signaling, increasing evidence suggests that the membrane itself plays an active role in both suppressing and enhancing signaling. Identifying and investigating this contribution has been challenging owing to the complex composition of the plasma membrane. We used cell-free expression to incorporate the epidermal growth factor receptor (EGFR) into nanodiscs with defined membrane compositions and characterized ligand-induced transmembrane conformational response and interactions with signaling partners using single-molecule and ensemble fluorescence assays. We observed that both the transmembrane conformational response and interactions with signaling partners are strongly lipid dependent, consistent with previous observations of electrostatic interactions between the anionic lipids and conserved basic residues near the membrane adjacent domain. Strikingly, the active conformation of EGFR and high levels of ATP binding were maintained regardless of ligand binding with high anionic lipid content typical of cancer cells, where EGFR signaling is enhanced. In contrast, the conformational response was suppressed in the presence of cholesterol, providing a mechanism for its known inhibitory effect on EGFR signaling. Our findings introduce a model of EGFR signaling in which the lipid environment can override ligand control, providing a biophysical basis for both robust EGFR activity in healthy cells and aberrant activity under pathological conditions. The membrane-adjacent protein sequence, likely responsible for the lipid dependence, is conserved among receptor tyrosine kinases, suggesting that active regulation by the plasma membrane may be a general feature of this important class of proteins.

## Introduction

The epidermal growth factor receptor (EGFR), the canonical receptor tyrosine kinase, maintains basic cellular processes in mammals.^1, 2^ EGFR phosphorylates adaptor proteins that trigger an array of signaling cascades responsible for cell proliferation and differentiation or, upon aberrant activity, leads to disorders such as cancer and fibrosis.^3, 4^ Ligand binding to the extracellular region of the receptor initiates a signaling response that propagates across the plasma membrane.^5^ The composition of the membrane changes from healthy to disease states in a manner known to influence EGFR activity.^6–9^ A basic understanding of the ligand-induced structural reorganization of the receptor has been established,^10–18^ yet the contribution of the plasma membrane in regulating this reorganization remains underinvestigated.^19^

The chemical composition of the human plasma membrane is complex. The membrane composition is well maintained in healthy cells, yet becomes dysregulated in a diseased state, where EGFR signaling is often aberrant.^20, 21^ In healthy cells, the plasma membrane is an asymmetric lipid bilayer containing 30% anionic lipids in the inner leaflet.^22^ These anionic lipids modulate the surface charge of the membrane, affecting protein localization and clustering.^23^ Anionic lipids also act as signaling molecules and regulate several fundamental cellular processes.^24–28^ The anionic lipid content increases in cancer cells, where EGFR signaling is typically increased.^29–32^ Similarly, dysregulation of lipid metabolism and distribution coupled with hyperactivation of EGFR are major hallmarks of neurodegenerative disorders, including Alzheimer’s disease, Parkinson’s disease, dementia, and sclerosis.^33, 34^ Cholesterol is the major sterol component of mammalian cell membranes, averaging 20% – 25% of the lipid bilayer.^35^ Cholesterol maintains the structural integrity and regulates the fluidity of the bilayer.^36, 37^ Increasingly, cholesterol has been implicated in the modulation of signal transduction and cellular trafficking.^38–40^ In the case of EGFR, cholesterol has been shown to suppress ligand-induced signaling.^41^ While a relationship between the plasma membrane and the signaling cascade has been established, how individual components, particularly anionic lipids and cholesterol, influence the protein-level transmembrane conformational response of EGFR remains unclear.

Several studies have investigated the role of anionic lipids and cholesterol for individual domains of EGFR. The plasma membrane was found to abrogate the catalytic activity of the intracellular kinase domain^42^ and reduce the ligand binding affinity for an unliganded protomer in a dimer of the extracellular domain, leading to a negative cooperativity model.^43–46^ Cholesterol has also been shown to impede phosphorylation for EGFR in vitro.^47^ In transmembrane-juxtamembrane constructs of EGFR, electrostatic interactions between the positively charged juxtamembrane region and the negatively charged anionic lipids have been identified.^48–53^ While these studies have provided some understanding of the impact of the plasma membrane on individual domains, transmembrane conformational signaling requires propagation through these domains to cross the membrane. Piecewise studies inherently cannot probe how such propagation is regulated by membrane composition.

Here, we report the observation of a transmembrane conformational response of EGFR in membranes with different compositions using single-molecule Förster resonance energy transfer (smFRET), ensemble fluorescence assays, and molecular dynamics simulations. These measurements showed a robust ligand-induced conformational response in membranes that mimic healthy cells, consistent with previous observations.^15, 18^ However, in membranes enriched with anionic lipids and/or cholesterol, we found a suppression of this response—revealing a previously unrecognized effect of these lipid components. These results indicate that membrane composition actively modulates EGFR function through both its charge and mechanics, playing a key role in robust signaling. Therapeutic development targeting EGFR may, therefore, require the context of the plasma membrane for optimal efficacy.

## Results and Discussion

### EGFR in membrane nanodiscs

EGFR-containing discoidal membranes, termed ‘nanodiscs’, were produced by in vitro co-expression of EGFR and an apolipoprotein in the presence of lipid vesicles, leading to self-assembly of the structures shown in Fig. 1a (Supplementary Fig. 2, 3).^54, 55^ Nanodiscs were produced with different ratios of 1-palmitoyl-2-oleoyl-sn-glycero-3-phosphocholine (POPC), 1-palmitoyl-2-oleoyl-sn-glycero-3-phospho-L-serine (POPS) and cholesterol and dimyristoylphosphatidylcholine (DMPC) (Supplementary Fig. 4, 5). POPC is a zwitterionic phospholipid forming neutral membranes, whereas POPS carries a net negative charge and provides anionic character to the bilayer.^56^ Both PC and PS lipids are common constituents of mammalian plasma membranes, with PC enriched in the outer leaflet and PS in the inner leaflet.^22^ Nine lipid environments with different combinations of anionic lipid and/or cholesterol content were compared: 0%, 15%, 30%, 60% POPS in POPC; 7.5%, 20% cholesterol in POPC and in 30% POPS in POPC; and DMPC, the fully saturated analog of the monounsaturated POPC. Zeta potential analysis was used to confirm the incorporation of POPS into the nanodiscs (Fig. 1b, Supplementary Fig. 6).^56^ Laurdan, a fluorescent membrane marker for membrane fluidity, was used to confirm the incorporation of cholesterol (Fig. 1c, Supplementary Fig. 7).^57^

**Figure 1.**
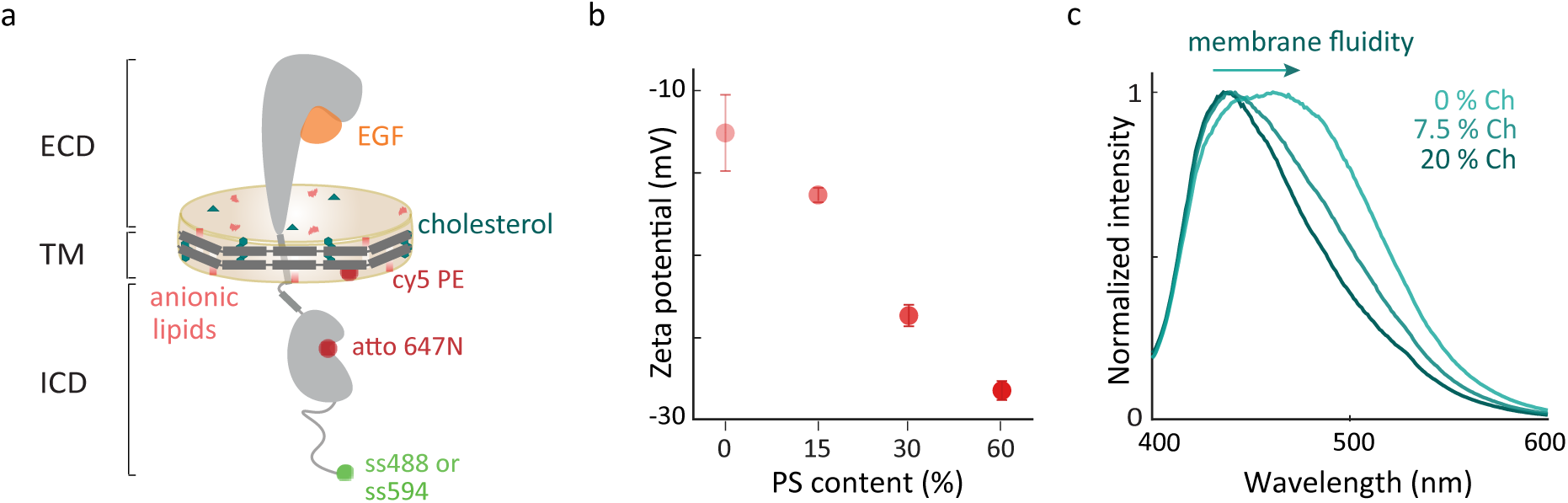
EGFR in different membrane environments. (a) Full-length EGFR (gray) embedded in a nanodisc. The nanodisc is a lipid bilayer (beige) belted by an amphiphilic apolipoprotein (dark gray). EGFR consists of a 618-amino acid extracellular region that binds EGF (orange), a 27-amino acid transmembrane-spanning domain, and an intracellular region, which is a 37-amino acid juxtamembrane domain, a 273-amino acid kinase domain and a 231-amino acid disordered C-terminal tail. Green and maroon spheres indicate the donor and acceptor dyes, respectively (Supplementary Fig. 1). (b) Mean of zeta potential distributions for EGFR in nanodiscs containing increasing amounts of anionic lipids (0%, 15%, 30% and 60% POPS). Error bars are from three technical replicates. (c) Ensemble fluorescence emission spectra (*λ_exc_*= 385 nm) of EGFR embedded Laurdan containing nanodiscs with increasing cholesterol.

A fluorescent donor-acceptor dye pair was incorporated into the nanodisc construct for fluorescence measurements. The donor dye (snap surface 488 or 594) was covalently attached to the snap tag at the EGFR C-terminus and the acceptor dye was introduced as a Cy5 labeled lipid at low concentration in the lipid bilayer or bound as Atto647N-labelled 1-ATP (Supplementary Fig. 8).^18, 55^ For the Cy5 labeled lipid, the steady state absorption and emission spectra, and lifetime of the acceptor dye were consistent between different nanodisc membrane compositions, indicating no lipid-dependent photophysics (Supplementary Fig. 9). The functionality of labeled EGFR in the nanodiscs was evaluated by Western blotting for phosphorylation of the tyrosine residues, which showed levels consistent with previously published assays on similar preparations (Supplementary Fig. 2d).^18, 54, 55^

### Role of anionic lipids in EGFR kinase activity

The enzymatic activity of the catalytic site in the EGFR kinase domain can be regulated through its accessibility to ATP and other substrates.^58^ To investigate how the membrane composition impacts accessibility, we measured ATP binding levels for EGFR in membranes with different anionic lipid content. 1 µM of fluorescently-labeled ATP analogue, atto647N-1 ATP, which binds irreversibly to the active site, was added to samples of EGFR nanodiscs with 0%, 15%, 30% or 60% anionic lipid content in the absence or presence of EGF. The fluorescence intensity from the bound ATP analogue and the fluorescence intensity from snap surface 488, which binds stoichiometrically to the snap tag at the EGFR C-terminus, were measured for each sample from a fluorescent gel image. The relative amount of ATP binding was quantified for each sample by normalizing to the EGFR content (Fig. 2a, b).

**Figure 2.**
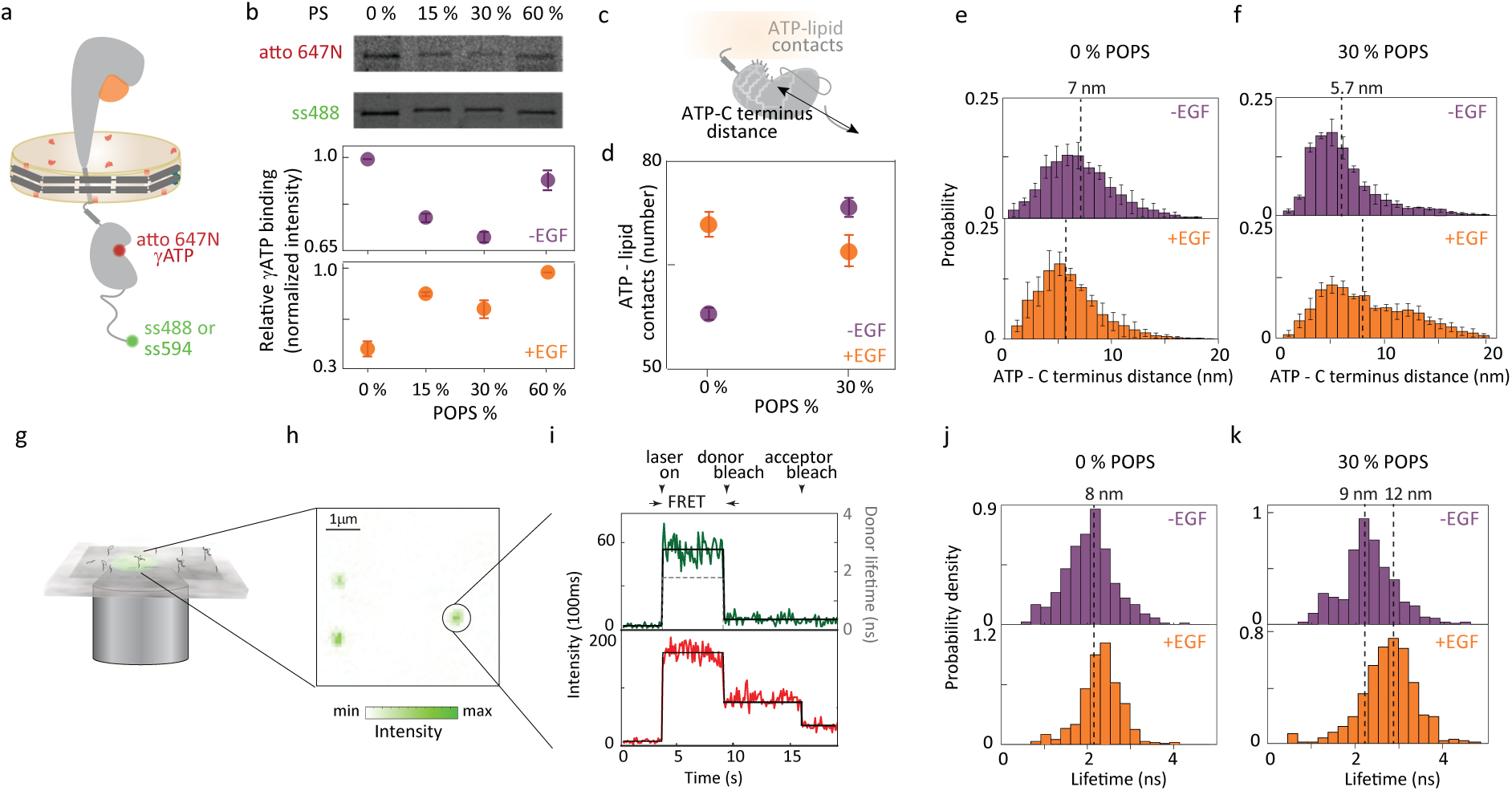
Membrane composition influences EGFR function through ATP binding. (a) Full-length EGFR in nanodiscs with atto647N γ ATP (red sphere) and snap surface 488 or snap surface 594 (green sphere). (b) Extent of ATP binding in different anionic environments quantified using the intensity of atto 647N (top) band normalized by the amount of EGFR produced as extracted from the intensity of ss488 band (center) as a function of negatively charged POPS lipids (bottom) in the absence (purple) and presence (orange) of EGF. Error bars from three independent biological replicates. (c) EGFR intracellular domain indicating ATP binding site-C-terminus distance and ATP-lipid contacts measured from molecular dynamics simulations. (d) Accessibility of ATP binding site quantified through the contact number between the ATP binding site and lipids in the absence (purple) and presence (orange) of EGF. Probability distributions of the distance between residue 721, the closest residue to the ATP binding site,^60^ and EGFR C-terminus for (e) neutral (0% POPS) and (f) 30% anionic lipids (30% POPS) without EGF (top); with 1 *µ*M EGF (bottom). Dotted lines indicate the medians on all histograms with corresponding distances on upper x-axis. (g) Schematic of multiparametric single-molecule confocal microscope. (h) Fluorescence intensity for a representative image (*λ_exc_* = 550 nm) where green spots are immobilized EGFR nanodiscs. (i) Representative fluorescence time trace from single-molecule FRET experiments showing number of detected photons for each 100 ms interval as intensity traces (green for donor; red for acceptor) with the average for each period of constant intensity (black solid line) and the corresponding donor lifetime (black dashed line). smFRET donor lifetime distributions with atto647N *I*ATP as acceptor and snap surface 594 as donor in (j) neutral (0% POPS) and (k) 30% anionic lipids (30% POPS) without EGF (top); with 1 *µ*M EGF (bottom).

How the level of ATP binding changed with anionic lipid content depended on the presence of EGF. In the absence of EGF, ATP binding was high in neutral bilayers (0% POPS) and in highly anionic (60% POPS) bilayers, but low at physiologically relevant POPS levels (15% and 30% POPS). This suggests that nucleotide binding is suppressed in the physiological regime, likely to ensure EGF can promote catalytic activity. In the presence of EGF, ATP binding overall increased with anionic lipid content with the highest levels observed in 60% POPS bilayers. In the neutral bilayer, ligand seemed to suppress ATP binding, indicating anionic lipids are required for the regulated activation of EGFR. Similar, and relatively high, levels of ATP binding were observed in the physiological regime, consistent with the model of EGF-promoted activity. The high ATP binding at 60% POPS —even without EGF —is consistent with the enhanced levels of catalytic activity characteristic of cancer cells, which show high anionic lipid content.^29–32^ In previous work, experiments and molecular dynamics simulations identified electrostatic interactions between anionic lipids and the kinase domain of EGFR.^15, 59^ These results highlight that membrane composition, through electrostatic interactions with anionic lipids, can regulate accessibility of the ATP binding site, likely as a mechanism to modulate EGFR catalytic activity.

The accessibility of the ATP binding site was also examined through additional molecular dynamics simulations (Fig. 2c-f). Simulations were carried out on EGFR in a neutral and partially (30%) anionic lipid bilayer in the absence and presence of EGF. For each condition, the number of contacts between the ATP binding site and the membrane was determined (Fig. 2c, d). The number of contacts was defined as the number of coarse-grained atom pairs between the lipid membrane and the ATP binding site that have a smaller than 16 Å distance. In the absence of EGF, increasing the anionic lipid content from 0% POPS to 30% POPS increased the number of ATP-lipid contacts from 58.6 *±* 0.7 to 74.4 *±* 1.2, indicating reduced accessibility, consistent with the experimental results and suggesting anionic lipids are required for ligand-induced EGFR activity. In the presence of EGF, increasing the anionic lipid content decreased the number of contacts from 71.8 *±* 1.8 to 67.8 *±* 2.4, indicating increased accessibility, again in line with the experimental findings. Because detection of EGFR relies on labeling at the C-terminus and ATP binding requires an intact kinase domain, the ATP-binding assay is for receptors that are properly folded and competent for nucleotide binding. The consistency between experimental results and MD simulations suggests that the observed lipid-dependent changes are more likely due to modulation of functional EGFR than to artifacts from misfolding.

### Anionic lipid dependence of kinase domain position

To experimentally investigate the intracellular conformations responsible for the lipid-dependent ATP accessibility, we performed smFRET measurements that probed the relative organization of the kinase domain and the C-terminal tail. EGFR was embedded in 0% and 30% POPS nanodiscs with snap surface 594 on the C-terminal tail as the donor and a fluorescently labeled ATP analog (atto647N) as the acceptor (Fig. 2a). The affinity tag on the belting protein was used for immobilization of EGFR nanodiscs at dilute concentration on a coverslip so the donor fluorescence could be recorded for individual constructs (Fig. 2g-i, Supplementary Fig. 10).^18^ These measurements monitored the donor fluorescence lifetime, which decreases with donor-acceptor distance or as energy transfer to the acceptor increases.^61^ The donor lifetime distributions of the EGFR samples in the absence and presence of EGF (1 µM) are shown in Fig. 2j, k, Table 1, Supplementary Table 1, 2). The corresponding donor-acceptor distances were calculated using the lifetime of the donor only sample as a reference (Supplementary Fig. 11). In the neutral bilayer (0% POPS), the distributions in the absence of EGF peaks at 8.1 nm (95% CI: 8.0–8.2 nm) and in the presence of EGF peaks at 8.6 nm (95% CI: 8.5–8.7 nm) (Table 1, Supplementary Table 1). In the physiological regime of 30% POPS nanodiscs, the peak of the donor lifetime distribution shifts from 9.1 nm (95% CI: 8.9–9.2 nm) in the absence of EGF 11.6 nm (95% CI: 11.1–12.6 nm) in the presence of EGF (Table 1, Supplementary Table 1), which is a larger EGF-induced conformational response than in neutral lipids. The larger conformational response observed in the presence of anionic lipids suggests that these lipids enhance the responsiveness of the intracellular domains to EGF, potentially facilitating interactions between C-terminal sites and adaptor proteins during downstream signaling.

**Table 1.**
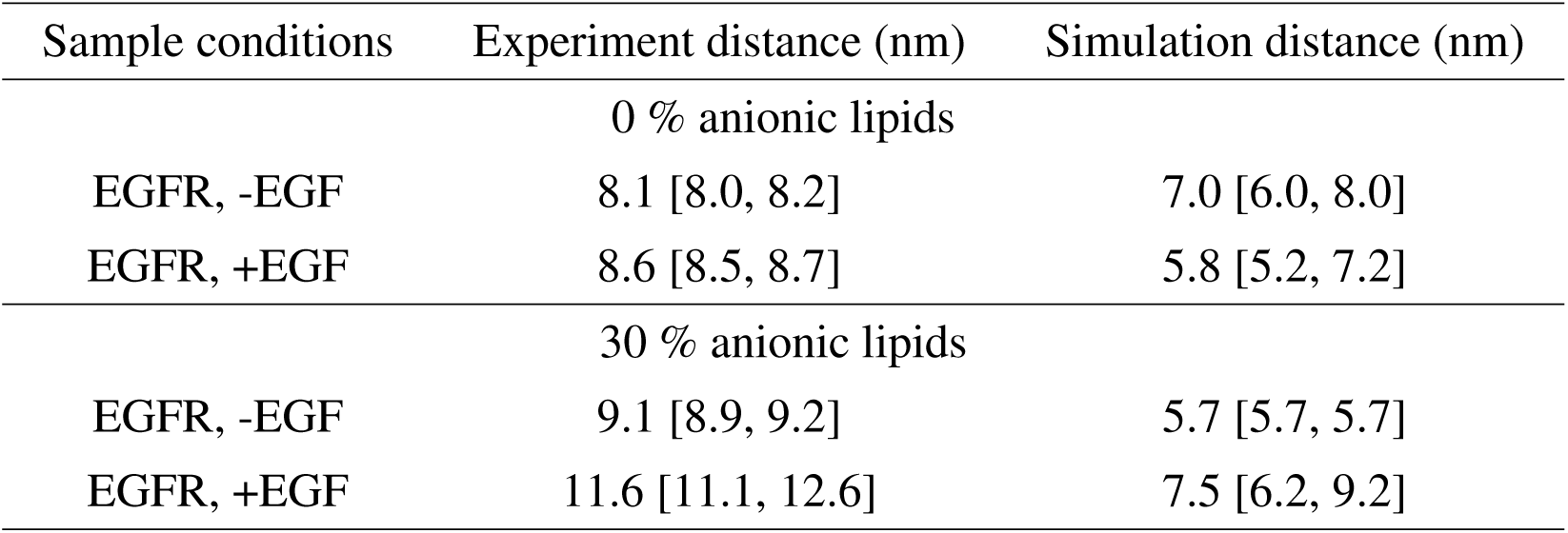
Median distances between the ATP binding site (residue 721) and the C-terminal end of EGFR from smFRET experiments and simulations. The distance values were extracted as the medians of the distributions in Figs. 2e, f, j, k in the main text. The numbers in parenthesis indicates the 95 % confidence interval for experiments and the minimal and maximal median value from three equal partitions of data for simulations. Ro = 7.5 nm for snap surface 594 and atto 647N

The same distance between C-terminus of the protein and ATP binding site was extracted from the molecular dynamics simulations for both neutral and 30% anionic membranes (Fig. 2c). In the neutral bilayer, the distance was 7.0 nm and 5.8 nm in the absence and presence of EGF, respectively (Fig. 2e, Table 1). While a small (∼1 nm) compaction was observed, both values correspond to an overall similar conformation. Upon introduction of 30% anionic lipids in the bilayer, the measured distance shifted from 5.7 nm in the absence of EGF to 7.5 nm in the presence of EGF (Fig. 2f, Table 1). Both experimental and computational results show a larger EGF-induced shift in the partially anionic bilayer, consistent with the notion that a partially anionic lipid bilayer provides a more native environment that supports proper receptor activation, compared to the non-physiological neutral membrane.

### smFRET measurements of EGFR in membrane nanodiscs

To further map out the conformational response, the overall intracellular conformation of EGFR in the absence and presence of extracellular EGF binding was captured using an additional series of smFRET measurements for different membrane compositions. EGFR nanodiscs containing 0%, 15%, 30% or 60% of the anionic lipid POPS doped in to the POPC bilayer were prepared with snap surface 594 on the C-terminal tail as the donor and the fluorescently-labeled lipid (Cy5) as the acceptor (Fig. 3a, Supplementary Fig. 2, 12). The donor lifetime was measured for all membrane compositions in the presence and absence of saturating concentrations (1 µM) of EGF. Histograms of the donor lifetimes were constructed for all conditions (Fig. 3, Supplementary Fig. 13, Table 2, Supplementary Table 3, 4).

**Figure 3.**
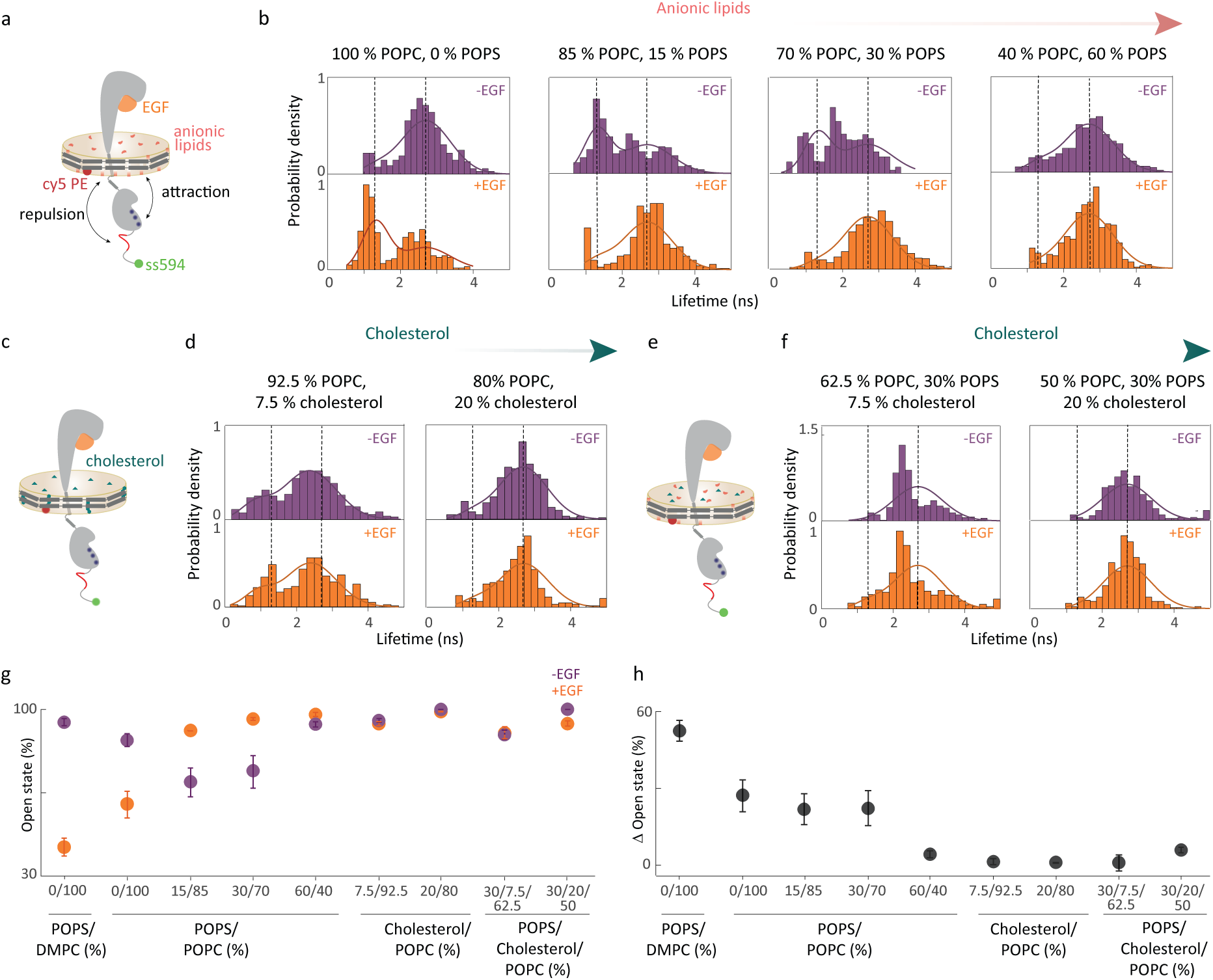
EGFR intracellular conformations depend on membrane composition of the nanodisc. (a) Full-length EGFR in nanodiscs with partially anionic lipids (red). The negatively charged residues on the C-terminal tail are indicated in red and the positively charged residues on the kinase domain are indicated in blue. smFRET donor fluorescence lifetime distributions in (b) 100% POPC, 0% POPS; 85% POPC, 15% POPS; 70% POPC, 30% POPS; 40% POPC, 60% POPS without EGF (top); with 1 *µ*M EGF (bottom). (c) Full-length EGFR in nanodiscs with cholesterol (teal). (d) smFRET donor fluorescence lifetime distributions in 92.5% POPC, 7.5% cholesterol; 80% POPC, 20% cholesterol without EGF (top); with 1 *µ*M EGF (bottom). (e) Full-length EGFR in nanodiscs with cholesterol and anionic lipids. (f) smFRET donor fluorescence lifetime distributions in 62.5% POPC, 30% POPS, 7.5% cholesterol; 50% POPC, 30% POPS, 20% cholesterol without EGF (top); with 1 *µ*M EGF (bottom). Dotted lines indicate the maxima from a global fit of all lifetime distributions to a double Gaussian distribution model using maximum likelihood estimation. The maxima correspond to a compact and an open conformation with a distance of 8 nm and 12 nm, respectively, between the EGFR C-terminal tail and the membrane bilayer. (g) The amplitude of the open conformation (in %) in all the eight different membrane compositions in the absence (purple) and presence of EGF (orange). (h) The change in amplitude induced by EGF (black). The amplitude change upon EGF addition is high (22% – 55%) in 0% – 30% POPS but reduces drastically (0% – 6%) upon introduction of cholesterol in the lipid bilayer. The error bars in (g) and (h) are from the global fit.

**Table 2.** Median distances between the membrane and C-terminal end of the protein from smFRET experiments. The distance values were extracted from the distributions shown in Fig. 3b in the main text. The numbers in parenthesis indicates the 95 % confidence interval for experiments. Asterisk (*) indicates distance was beyond the FRET range for the snap surface 594 and cy5 dye pair (Ro = 8.4 nm).^61^

| Sample conditions | Distance (nm) |
| --- | --- |
| 0 % POPS, 100 % POPC; -EGF | 12.1 [11.6, 12.7] |
| 0 % POPS, 100 % POPC; +EGF | 8.8 [8.2, 9.6] |
| 15 % POPS, 85 % POPC; -EGF | 9.2 [8.8, 9.6] |
| 15 % POPS, 85 % POPC; +EGF | 11.9 [11.4, 12.4] |
| 30 % POPS, 70 % POPC; -EGF | 9.2 [9.1, 9.3] |
| 30 % POPS, 70 % POPC; +EGF | *13.9 [12.5, 14.4] |
| 60 % POPS, 40 % POPC; -EGF | 11.4 [11.1, 11.7] |
| 60 % POPS, 40 % POPC; +EGF | 11.2 [10.9, 11.6] |

Examination of the distributions showed signatures of a bimodal structure, indicating two-state behavior with a distinct conformational equilibrium for each sample (Supplementary Table 5). To quantify the equilibrium for the samples, we globally fit the lifetime distributions for all the samples using maximum likelihood estimation with a double Gaussian distribution model.^62^ In global fit, the peak positions and widths are shared parameters between the samples and only the relative amplitudes of the two states changes for each distribution. The results of the global fitting are displayed as solid lines in Fig. 3b, d, f, Supplementary Table 6. The global fitting identified peaks at 1.3 ns (∼8 nm) and 2.7 ns (∼12 nm) with widths of 0.35 ns (∼1.9 nm) and 0.67 ns (∼3 nm), respectively. These two peaks represent a compact and an open conformation, *i.e.*, the C-terminal tail close to and away from the lipid bilayer surface, respectively.

### Anionic lipid dependence of transmembrane conformational response

To investigate the role of anionic lipids in EGFR transmembrane conformational response, the smFRET distributions were compared for nanodisc samples with increasing anionic lipid content (Fig. 3a, Table 2, Supplementary Table 3, 4). In the physiological regime for anionic lipid content (15%–30% POPS; Fig. 3b), the smFRET distributions exhibited a EGF-induced conformational response. In the absence of EGF, a conformational equilibrium was observed where the amplitude of the open conformation was similar at 69% and 74% for 15% and 30% POPS, respectively (Fig. 3b, Supplementary Table 6). In the presence of EGF, the conformational equilibrium was dominated by the open conformation with an amplitude of 91% and 96% for 15% and 30% POPS, respectively (Fig. 3b, Supplementary Table 6). For both lipid compositions, this corresponds to an increase in amplitude of the open conformation by ∼25%. (Fig. 3g, h). These results suggest that EGF-induced transmembrane conformational response is robust around the physiological anionic lipid content (Fig. 3g, h, Supplementary Table 6).

In the neutral bilayer (0% POPS; Fig. 3b), the distributions also exhibited a EGF-induced conformational response, but the response was the reverse of that in the physiological regime. As described in the previous section, in the absence of EGF, the conformational equilibrium was dominated by the open conformation at 87% whereas in the presence of EGF an equilibrium was observed where the amplitude of the open conformation was 60% (Fig. 3b, Supplementary Table 6). The reversal of the conformational response in the neutral bilayer strongly suggests that electrostatic interactions are playing an important role in transmembrane conformational response, consistent with previous smFRET experiments on EGFR.^18^

In the highly anionic bilayer (60% POPS; Fig. 3b), the EGF-induced conformational response nearly disappeared. In the absence of EGF, the conformational equilibrium was dominated by the open conformation > 90%, which, in the presence of EGF, increased slightly by ∼5%(Fig. 3b, g, h, Supplementary Table 6). The dominance of the open conformation in both conditions is similar to the EGF-bound conditions in the physiological regime, indicating that the receptor is held in the active conformation by the anionic bilayer. The amplitude of the open conformation correlated with the ATP binding site accessibility from the experimental and computational studies (Supplementary Fig. 14), consistent with a picture in which the open conformation enables access by all signaling partners. High levels of EGFR signaling are often present in cancer cells, where anionic lipid content is also high.^29^ Thus, the open conformation in the anionic bilayer may be constituitively active - and thereby the biophysical origin of the high signaling that drives cancer cell growth. Similar effects may be playing a role in the observed overactivation of EGFR in neurodegenerative disorders. Anionic lipids are increasingly recognized as biomarkers for neurodegenerative diseases, with EGFR emerging as a dual molecular target for both cancer and Alzheimer’s disease.^63^ Specifically, tyrosine kinase inhibitors, which are commonly used for cancer treatment, show promise in mitigating Alzheimer’s disease by targeting this overactive EGFR signaling pathway.^64^

### Electrostatic interactions in transmembrane conformational response

The molecular dynamics simulations were further examined to investigate how anionic lipids influence EGFR transmembrane coupling. EGF binding was found to switch the extracellular domain from prostrate on the membrane to upright,^15^ which, in turn, switched the transmembrane domain from a tilted to vertical orientation.^18^ The orientation switch caused the juxtamembrane domain to move from embedded inside to extended outside the lipid bilayer.^18^ The simulations showed the position of the juxtamembrane domain influenced two key interactions in the intracellular domain: (1) attraction between the basic residues in the juxtamembrane/kinase domains and the anionic lipids; and (2) repulsion between the acidic residues on the N-terminal portion of the C-terminal tail and the anionic lipids (Supplementary Fig. 1). The competition between these interactions dictates the overall conformation of the intracellular domain.

In the absence of EGF, the more embedded juxtamembrane domain meant that the attraction between the juxtamembrane/kinase domains and the lipids dominated, inducing a more closed conformation. The dominant closed conformation was consistent with the high compact state percentage observed in the smFRET distributions for 15% POPS (Fig. 3b). Within this picture of competing electrostatic interactions, the repulsion between the C-terminal tail and the anionic lipids is expected to dominate at high anionic lipid content likely because the number of negative charges on the C-terminus is greater than the number of positive charges on either the juxtamembrane or kinase domains. Consistently, a higher open state percentage was observed for 30% and 60% POPS. In the presence of EGF, the more extended juxtamembrane domain separated the juxtamembrane/kinase domains and the lipids, decreasing their attraction and leading to a more open conformation again consistent with the higher open state percentage for all partially anionic bilayers.

Oncogenic mutations such as K745E and K757E introduce negative charges into the kinase domain, which likely decrease the attraction between the kinase domain and the anionic lipids to allow the repulsion and thus the open conformation to dominate.^65, 66^ As the open conformation also dominates in the presence of EGF (Fig. 3b, g), these mutations likely increase signaling levels, consistent with their role in cancer. The electrostatic attraction between the kinase domain and anionic lipids described above was previously found to restrict the access of the tyrosines on the C-terminal tail to the catalytic site in the kinase domain.^59^ However, the electrostatic repulsion between the C-terminal tail and the lipids had not been captured due to the absence of the C-terminal tail in the previous work.^67^

### Cholesterol inhibits transmembrane conformational response

To investigate the mechanism behind the suppression of EGFR signaling by cholesterol, smFRET experiments were also performed on nanodisc samples with cholesterol incorporated into the lipid bilayer (Fig. 3c-f). The smFRET distributions were statistically the same in the presence and absence of EGF for all cholesterol-containing samples (Fig. 3d, f, Supplementary Figs. 15, 16, Table 3, Supplementary Table 7-12), revealing the suppression of the EGF-induced conformational response.

**Table 3.** Median distances between the membrane and C-terminal end of the protein from smFRET experiments. The distance values were extracted from the distributions shown in Fig. 3d and 3f in the main text. The numbers in parenthesis indicates the 95 % confidence interval for experiments. Asterisk (*) indicates distance was beyond the FRET range for the snap surface 594 and cy5 dye pair (Ro = 8.4 nm).^61^

| Sample conditions | Experiment distance (nm) |
| --- | --- |
| 92.5 % POPC, 7.5 % Cholesterol; -EGF | 12.1 [11.6, 12.6] |
| 92.5 % POPC, 7.5 % Cholesterol; +EGF | 12.2 [11.5, 13.2] |
| 80 % POPC, 20 % Cholesterol; -EGF | 11.1 [10.7, 11.5] |
| 80 % POPC, 20 % Cholesterol; +EGF | 10.9 [10.6, 11.2] |
| 62.5 % POPC, 30 % POPS, 7.5 % Cholesterol, -EGF | 10.4 [10.2, 10.5] |
| 62.5 % POPC, 30 % POPS, 7.5 % Cholesterol, +EGF | 10.3 [10.2, 10.4] |
| 50 % POPC, 30 % POPS, 20 % Cholesterol, -EGF | *12.9 [11.5, 12.3] |
| 50 % POPC, 30 % POPS, 20 % Cholesterol, +EGF | *13.0 [11.5, 12.1] |
| 100 % DMPC; -EGF | 11.4 [11.1, 11.6] |
| 100 % DMPC; +EGF | 8.1 [8.1, 8.2] |

In the physiological regime of cholesterol content (20%), the distributions were dominated by the open conformation with an amplitude of ∼95% (Fig. 3g, h, Supplementary Table 9, 12). For low cholesterol content (7.5%), the distributions were still dominated by the open conformation with a slightly smaller amplitude of ∼90%, corresponding to a shift in the peak of the distribution to shorter distances (Supplementary Table 9, 12). Again, the distributions were statistically the same in the presence and absence of EGF (Supplementary Figs. 15, 16, Supplementary Table 7, 10), indicating only a small amount (≤7.5%) of cholesterol is required to suppress the conformational response.

Consistent with the suppression of the conformational response, several previous observations show that cholesterol impedes EGFR function, leading to the hypothesis that it must be sequestered for proper signaling.^68^ An artificial increase in the cholesterol content of the plasma membrane for A431 cells or HEp-2 cells led to reduced ligand-induced EGFR activation.^41^ Cholesterol depletion by the drugs MpCD or U18666A also increased dimerization and phosphorylation of EGFR.^41, 69^ The introduction of cholesterol into the membrane of proteoliposomes containing reconstituted EGFR resulted in decreased kinase activity, which was attributed to a change in membrane fluidity.^47^ Although EGFR is locked in the EGF-bound configuration, the suppressed conformational response may play a role in the biophysical mechanism behind the ability of cholesterol to impede EGFR signaling.^41^ Remarkably, high anionic lipids and cholesterol content produce the same EGFR conformations but with opposite effects on signaling—suppression or enhancement. Both conditions, however, underscore the complexity and sensitivity of membrane regulation in cell signaling.

### Mechanism of cholesterol inhibition of EGFR transmembrane conformational response

Cholesterol is known to impact both the thickness and the fluidity of lipid bilayers. Previous computational and experimental results showed high cholesterol can increase the thickness of a phospholipid bilayer, defined as the separation between lipid phosphate head groups, by ∼0.3 – 0.5 nm.^70, 71^ Evidence suggests that transmembrane helices tilt to minimize hydrophobic mismatch,^71^ changing the conformation of embedded membrane proteins. The incorporation of cholesterol into the lipid bilayer has also been shown to alter its fluidity. The altered fluidity was also found to modulate the mobility and oligomerization of ErbB2, a member of the EGFR family.^72^ Either factor —bilayer thickness or fluidity —could, therefore, give rise to the suppressed conformational response observed in the smFRET measurements.

To identify the mechanistic origin of the cholesterol-induced suppression of the conformational response, smFRET experiments were performed on EGFR-containing nanodiscs with two different lipids, DMPC and POPC (Supplementary Figs. 17, Table 3, Supplementary Table 3-6). DMPC is a fully saturated version of POPC that forms a thinner bilayer (3.7 nm for DMPC and 3.9 nm for POPC,^73^ which is similar in magnitude to the thickness change observed upon the addition of cholesterol^70, 71^). The fluidity of the lipid bilayer is described through the order parameter (S), which is the average ordering of the lipid chains. S is ∼0.3 for nanodiscs of the same size and containing the same number of DMPC or POPC lipids, indicating comparable fluidity.^74^

In both POPC and DMPC bilayers, the distribution was dominated by the open conformation (∼12 nm) in the absence of EGF, whereas the equilibrium shifted towards the compact conformation (∼8 nm) in the presence of EGF (Supplementary Figs. 17). The amplitude of the open conformation decreased from ∼87% to 60% in its presence for POPC and decreased from 95% in the absence of EGF to 42% in its presence for DMPC (Supplementary Table 6). The similar behavior between POPC and DMPC strongly implies the transmembrane conformational response is independent of bilayer thickness, suggesting that the decreased membrane fluidity in the presence of cholesterol is likely responsible for the suppression of the conformational response.

### Implications of lipid dependence on EGFR and other membrane receptors

EGFR activation has long been understood through the lens of ligand-induced structural rear rangements and downstream protein-protein interactions. However, studies of these components were unable to evaluate the role of the surrounding membrane. Our findings introduce a direct effect of membrane composition on the conformational response of EGFR, explaining previous observations of membrane-mediated regulation of downstream signaling. Here, we report the discovery that the lipid composition—through both electrostatics and membrane mechanics–can override ligand-driven activation to regulate receptor function (Fig. 4).

**Figure 4.**
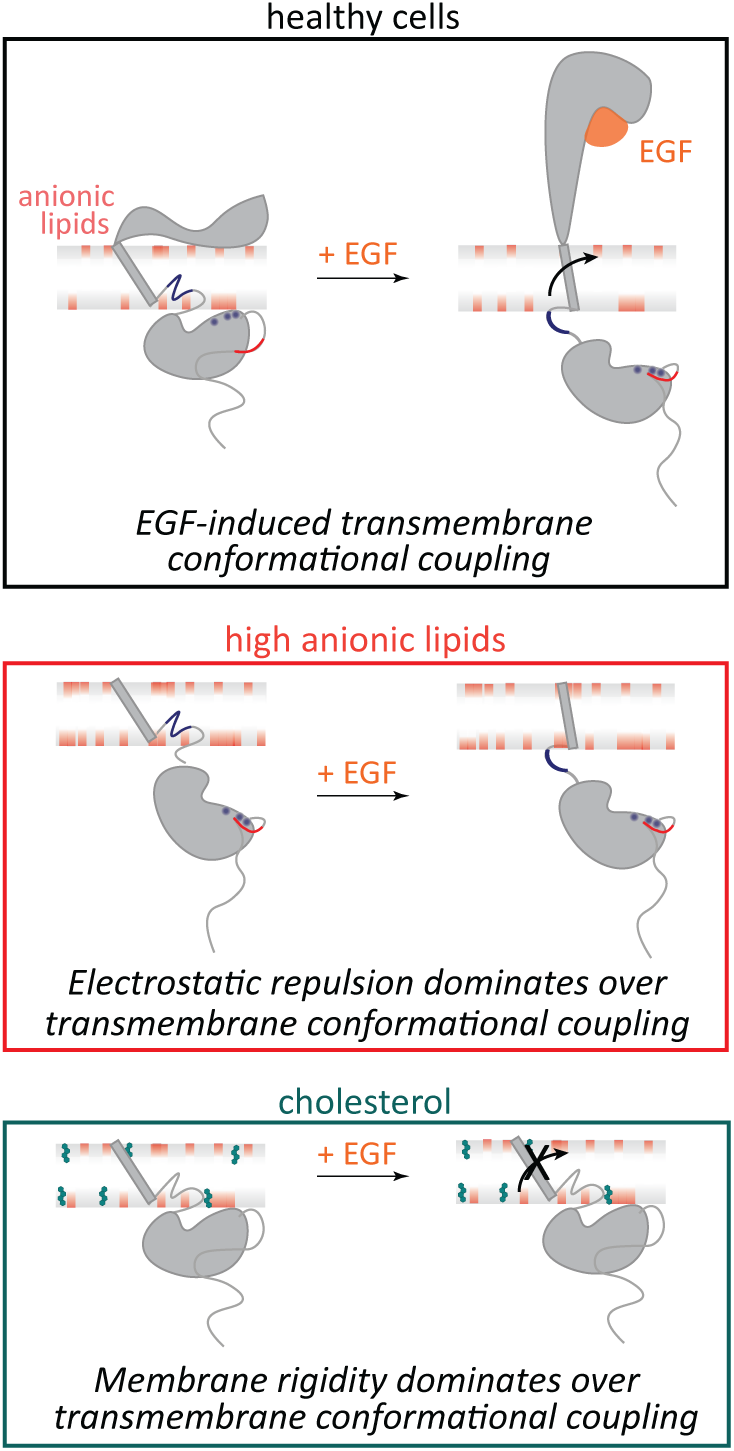
Membrane composition influences EGFR conformation and function. (Top) In healthy cells, EGF binding promotes transmembrane conformational coupling between the extracellular and intracellular domains of EGFR, enabling kinase activation through established signaling mechanisms. (Middle) Our results demonstrate that in membranes with high anionic lipid content, electrostatic repulsion between the negatively charged lipids and the kinase domain overrides EGF-induced transmembrane conformational coupling. (Bottom) Our results demonstrate that in cholesterol-rich membranes, increased membrane rigidity overrides EGF-induced transmembrane conformational coupling. These findings establish membrane composition as a dominant regulator of EGFR signaling, independent of EGF binding.

Notably, we found that the conformational response is suppressed in membrane compositions known to impede healthy signaling, *i.e.*, high cholesterol or high anionic lipid content. While autoinhibitive interactions of lipids with individual EGFR domains had been established previously,^15, 42, 59^ our results show that the protein-lipid interplay is crucial to maintain a ligand-induced transmembrane conformational response, which is the key regulator of healthy signaling. Our results complement and expand upon previous molecular dynamic simulations of the kinase domain that increasing amounts of anionic lipids (up to 30%) induce a corresponding decrease in the accessibility of the ATP binding site.^15^ Similar interactions were observed for PIP2 lipids as for PS,^18, 75–77^ suggesting similar effects may be induced during PIP2-mediated processes. Apart from phospholipids and sterols, glycolipids have been shown to have pronounced effect on EGFR activity.^78, 79^ Ganglioside GM3 inhibits cell motility via down regulation of ligand-stimulated EGFR phosphorylation and signaling.^80, 81^ Our results suggest that these components may function through conformational effects similar to the ones observed here.

In addition to EGFR, membrane composition is crucial for the activity of many membrane proteins.^82^ For example, rhodopsin has an important and functional interaction with G proteins that hinges on the presence of POPS lipids. These lipids are essential for maintaining the structure of its key amphipathic helix, similar to the POPS dependence observed here.^83, 84^ Ap-propriate lipid size is also necessary to match membrane thickness with transmembrane regions of integral proteins, preventing hydrophobic mismatches that could cause structural anomalies or significantly impair protein function,^85, 86^ such as 80% reduction in Ca^2+^-ATPase activity.^87, 88^ Furthermore, the saturation level of lipid hydrocarbon tails, influencing their melting temperature, is critical in determining membrane bilayer states and protein activity.^89^ Thus, the insights into the membrane dependence of EGFR signaling reported here could be universal to other membrane receptors, particularly the family of proteins that share the same structural homology and perform other significant functions and suggest that the membrane has broad implications in cellular signaling.

## Methods

### Production of labeled full-length EGFR nanodiscs

Fluorescently-labeled EGFR in nanodiscs was produced and characterized as described in previously published protocols.^18, 54, 55^ ApoA1△49 in pIVEX2.4d vector (Genscript) and full-length EGFR (1210 amino acids) with SNAP tag at the C-terminus in SNAP T7 vector (Genscript) was added to the cell-free reaction contents (Thermofisher Scientific) (Supplementary Fig. 2). Briefly, the E.Coli slyD lysate, in vitro protein synthesis E.Coli reaction buffer, amino acids (-Methionine), Methionine, T7 Enzyme, protease inhibitor cocktail (Thermofisher Scientific), RNAse inhibitor (Roche) and DNA plasmids (20ug of EGFR and 0.2ug of ApoA1△49) were mixed with different lipid mixtures. The DNA template ratio of EGFR:ApoA149 = 100:1 was empirically chosen by testing different ratios on SDS-PAGE gels and selecting the condition that maximized full-length EGFR expression in DMPC lipids (Supplementary Fig. 3). The lipid mixtures were made by sonicating different percentages of 1,2-dimyristoyl-sn-glycero-3-phosphocholine (DMPC) (Avanti Polar Lipids), 1-palmitoyl-2-oleoyl-sn-glycero-3-phosphocholine (POPC) (Avanti Polar Lipids), 1-palmitoyl-2-oleoyl-sn-glycero-3-phospho-L-serine (POPS) (Avanti Polar Lipids) and cholesterol (Sigma-Aldrich) keeping the total lipid concentration at 2mg/mL for the cell-free reaction mixture. For introducing a single acceptor into the nanodisc, a molar ratio of 500:1 lipid:cy5 labeled, 2-dioleoyl-sn-glycero-3-phosphoethanolamine (DOPE) lipids (Avanti Polar lipids) was added to the above solution after bath sonication. The above solution was incubated at 25*^o^*C, 300 rpm for 30 minutes. The mixture was supplemented with E.Coli feed buffer, amino acids (-Methionine) and Methionine (total volume of cell-free reaction is 250 µL) and incubated for additional 8 hours. 500 nM of snap surface 594 (New England Biolabs) was next added to the above mixture and incubated at 37*^o^*C, 300 rpm for 35 minutes. Snap surface 594, a derivative of Atto594 with benzylguanine functionality reacts in a near-stoichiometric efficiency with genetically-encoded snap tag.^90^

### Affinity Purification of labeled EGFR nanodiscs

500 µL of Ni-NTA resin slurry (Qiagen) was added to a 2 mL plastic column (Bio-Rad Laboratories). The resin was washed with double distilled water and equilibrated with 3 mL of native lysis buffer (50 mM NaH_2_PO_4_, 300 mM NaCl, pH 8.0). The cell-free reaction post labeling was added to 500 µL of lysis buffer on the incubated column and incubated at 4*^o^* C for 2 hours. After the flowthrough was collected, the column was washed with lysis buffer (3 ×1 mL) followed by lysis buffer containing 10 mM imidazole (2 ×1 mL) and lysis buffer containing 25 mM imidazole (2 ×1 mL) to remove all the non-specific interactions of the reaction mixture and free dye from the column. The EGFR nanodiscs were eluted with lysis buffer containing 400 mM (2 ×500 µL) imidazole. The eluted fractions were concentrated using 50 kDa, 500 µL spin filters (Sigma-Aldrich) by centrifugation.

### Protein content for labeled EGFR nanodiscs

SDS-PAGE was used to confirm the production of both belt protein (at 25 kDa) and EGFR (at 160 kDa). Samples were mixed with 2×Laemmli sample buffer (Bio-Rad Laboratories), 2.5 % 2-mercaptoethanol (Sigma-Aldrich) and boiled for 5 minutes at 95*^o^* C before running on precast stain-free gels from Bio-Rad Laboratories. Precision Plus Protein unstained standard (Bio-Rad Laboratories) marker was used for the stainfree imaging and Prestained NIR protein ladder (ThermoFisher Scientific) for fluorescence imaging. Gels were run at 170 V for 45 minutes. Stain-free and fluorescent imaging was performed on ChemiDoc Imaging System (Bio-Rad Laboratories). The other proteins appearing in the stainfree gel are proteins not expressed completely during the cell-free reaction or the transcription and translation machinery of the cell-free reaction. The specificity of snap surface 594 fluorophore binding to EGFR was confirmed through the fluorescence gel.

### Transmission Electron Microscopy

5 µL of cell-free expressed EGFR nanodiscs in 1×PBS buffer (137 mM NaCl, 2.7 mM KCl, 10 mM Na_2_HPO_4_, 1.8 mM NaH_2_PO_4_, pH 7.4) was added to glow-discharged carbon coated 400 mesh copper grids (Electron Microscopy Sciences) and incubated for 5 minutes at room temperature to allow nonspecific binding of the nanodiscs to the grids. The solutions were removed by gently blotting the side of the grid with filter paper. The grids were subsequently incubated with 5 µL of 2 % aqueous uranyl acetate for 30 seconds. Excess stain was removed similarly to the sample. The grids were air dried, and then imaged on a FEI Tecnai transmission electron microscope (120 kV, 0.35 nm point resolution). The distribution of disc sizes was analyzed using Image J software.

### Dynamic light scattering

The EGFR nanodiscs in 1×PBS buffer were filtered using 0.2 µm syringe filters and their dynamic light scattering measurements were performed on a DynaPro NanoStar (Wyatt Technologies, USA). Each measurement represents an average of 50 individual runs. Substantial heterogeneity in nanodisc lipid composition, such as uneven incorporation of cholesterol, would be expected to broaden the DLS distributions. However, comparison of the full width at half maximum (FWHM) values from the DLS distributions showed no significant broadening between cholesterol-containing and cholesterol-free nanodiscs (Mann-Whitney U test, p = 0.486; n = 4 for each group).

### Zeta potential measurements to quantify surface charge of nanodiscs

Titrations (0 %, 15 %, 30 % and 60 %) of negatively charged POPS lipids in neutral POPC lipids were performed to determine the surface charge of the nanodiscs with increasing negatively charged lipid content. Zeta potential measurements were performed on a Malvern Zetasizer Nano -– ZS90 (Malvern, UK), with a backscattering detection at a constant 173*^o^* scattering angle, equipped with a laser at 4mW, 633 nm. Dip cell ZEN1002 (Malvern UK) was used in the zeta-potential experiments. EGFR loaded nanodiscs produced from cell-free reactions were purified as mentioned above and buffered exchanged to 0.1 ×PBS. A final volume of 650 uL was prepared and transferred into the zeta dip cell. For each sample, a total of five scans, 30 runs each, with an initial equilibration time of 5 minutes, were recorded. All experiments were performed at 25*^o^* C.Values of the viscosity and refractive index were set at 0.8878 cP and 1.330, respectively. Data analysis was processed using the instrumental Malvern’s DTS software to obtain the mean zeta-potential value. This ensemble measurement reports the average surface charge of the nanodisc population, verifying incorporation of anionic POPS lipids.

### Fluoresence measurements with Laurdan to confirm cholesterol insertion into nanodiscs

Vesicles were prepared by mixing phospholipids and cholesterol in the desired ratios in chloroform such that the total concentration of the lipids was 20mg/mL. Laurdan was added in 100:1 lipid:laurdan ratio. The solvent was first evaporated by nitrogen flow followed by overnight drying in a vacuum desiccator. The dried samples were resuspended in water such that the total concentration of the lipids was 20mg/mL. The samples were then heated to 70°C and vortexed for 5-6 hours. After vortexing, the lipids were used in the cell-free reaction as described above. The EGFR nanodiscs containing laurdan were purified as described above and diluted with 1X PBS. The fluorescence excitation and emission spectra were recorded for the EGFR nanodiscs containing 0%, 7.5% and 20% cholesterol in the nanodiscs in Cary Eclipse fluorescence spectrophotometer (Agilent). The excitation spectrum was recorded by collecting the emission at 440 nm and emission spectra was recorded by exciting the sample at 385 nm. Laurdan fluorescence provides an ensemble readout of membrane order and confirms cholesterol incorporation into the nanodisc population. While laurdan does not resolve the composition of individual nanodiscs, prior work has shown that POPC-cholesterol mixtures are miscible without forming cholesterol-rich domains,^91, 92^ thus the observed ordering changes likely reflect the intended input cholesterol content at the ensemble level.

### Phosphorylation of EGFR in nanodiscs

Western Blot was performed using the Trans-Blot Turbo Transfer System (Bio-Rad Laboratories). The pre-loaded program for high molecular weight protein transfer was used for membrane transfer. After the transfer, the membrane was blocked in 5 % Non Fat Dry Milk (prepared in TBST buffer) for the anti-EGFR Western blots, and in 5 % BSA (prepared in TBST buffer) for the anti-phosphotyrosine Western blots for 20 minutes at room temperature. The membrane was then incubated in primary antibody overnight at 4*^o^* C. Following day, the membrane was washed and incubated with secondary antibody for 1 hour at room temperature. The primary antibodies, secondary antibodies and dilutions are listed in Supplementary Table 13. The fluorescence band detection was done using the ChemiDoc Imaging System (Bio-Rad Laboratories).

### Preparation of EGF ligand

Human EGF produced in E. Coli was purchased from Gold Biotechnology (Catalog Number: 1150-04-100). 1 µM EGF was prepared in 1×PBS buffer.

### ATP binding experiments

Full-length EGFR in different lipid environments was prepared using cell-free expression as described above. 1µM of snap surface 488 (New England Biolabs) and atto647N labeled gamma ATP (Jena Bioscience) was added after cell-free expression reaction and incubated at 30*^o^*, 300 rpm for 60 minutes. 1µM of atto647N gamma ATP was used, corresponding to a concentration near the reported K*_m_* of 5.2 µM for ATP binding to the isolated EGFR kinase domain,^93^ ensuring sensitivity to lipid-dependent changes in ATP accessibility. Purification using Ni-NTA affinity was performed and the samples were concentrated as described above. The samples were run on a SDS-PAGE gel at 170V for 40 minutes and imaged using the ChemiDoc Imaging System (Bio-Rad Laboratories).

### Fluorescence Spectroscopy

The His-tag present on the belt protein (ApoA1△49) was used to immobilize the EGFR-nanodisc constructs onto the microscope coverslip via Ni-NTA affinity. The purified EGFR nanodiscs were diltued to ∼500 pM in 1×PBS buffer and incubated for 15 minutes on the Ni-NTA coated glass (from Microsurfaces, Inc.) and flushed with solution containing 2 mM 6-hydroxy-2,5,7,8-tetramethylchroman-2-carboxylic acid (Sigma-Aldrich), 25 nM protocatechuate-3,4-dioxygenase (Sigma-Aldrich) and 2.5 mM protocatechuic acid (Sigma-Aldrich). Fluorescence experiments were then carried out on a home built confocal microscope.^94^ A Ti-Sapphire laser (Vitara-S, Coherent: λ*_c_* = 800 nm, 70 nm bandwidth, 20 fs pulse duration, 80 MHz repetition rate) was focused into a non-linear photonic crystal fiber (FemtoWhite 800, NKT Photonics) to generate a supercontinuum. Excitation light was then spectrally filtered for pulses centered at 550 nm or 640 nm and focused with an oil immersion objective lens (UPLSAPO100×, Olympus, NA = 1.4). Fluorescence emission was collected by the same objective and fed to the avalanche photodiodes (SPCMAQRH-15, Excelitas). A 5 µm ×5 µm area of a coverslip with immobilized receptors was scanned. Diffraction limited and spatially separated single molecule spots were then probed individually by unblocking the laser beam to record fluorescence until photo-bleaching. For 550 nm excitation, fluorescence was separated with a dichroic filter SP01-561RU (Laser 2000) and passed through FF01-629/56-25 (Semrock) for donor fluorescence collection. For experiments at 640 nm, ET 645/30×(Chroma) was used as the excitation filter, FF01-629/56-25 (Semrock) as the dichroic and FF02-685/40-25 (Semrock) for acceptor fluorescence collection. The laser power for the experiments was 2-3 µW at the sample plane.

Florescence emission was binned at 100-ms resolution to generate fluorescence intensity traces for both the donor and acceptor channels. Traces with a single photobleaching step for the donor and acceptor were considered for further analysis. Regions of constant intensity in the traces were identified by a change-point algorithm.^95^ Donor traces were assigned as FRET levels until acceptor photobleaching. The presence of empty nanodiscs does not influence these measurements, as photobleaching and single-molecule FRET analyses selectively report on receptor-containing nanodiscs. Consecutive bunches of 1000 photons in the donor channel were used to construct fluorescence decay curves for the FRET levels.^96^ The photons were histogrammed and the distributions were fit to a mono-exponential function convolved with the instrument response function (IRF) and summed with a separately-measured background term. The fit was performed using a Maximum Likelihood Estimator (MLE), which has been shown to be more accurate in the single-molecule regime.^94, 97^ The extracted lifetimes were used to construct histograms with bin sizes estimated from the square root of the total number of photon bunches.

The donor-acceptor distance (r in nm) was estimated using the following relation:^61, 98^

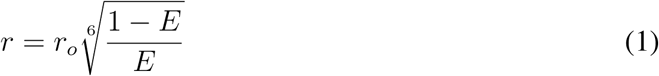

where r*_o_*is the calculated Förster distance (8.4 nm for snap surface 594 & cy5 dye pair; 7.5 nm for snap surface 594 & atto 647N dye pair)^61^ and the FRET efficiency (E) being experimentally measured as:

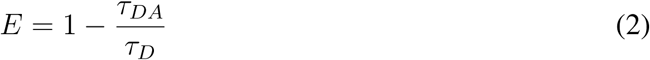

τ*_DA_* is the fluorescence lifetime of the donor in the presence of an acceptor, and τ*_D_* is the lifetime of donor-only construct. The distance between the donor and acceptor was quantified using a reference lifetime determined with a separately-characterized donor-only construct (Supplementary Figs. 11, 13, 15-17). For the smFRET measurements on the labeled constructs, the Cy5-lipid is confined to one side of the membrane for the duration of the measurement,^99^ and so the extracted distances are an average over the rapid translational diffusion of the labeled dye across the surface of the membrane nanodisc.^100^

### Model Selection and Statistical Analysis

Global fitting of lifetime distributions was performed across all experimental conditions using maximum likelihood estimation. Both two-Gaussian and three-Gaussian distribution models were evaluated as described previously.^62^ Model performance was compared using the Bayesian Information Criterion (BIC), ^101^ which balances model likelihood and complexity according to

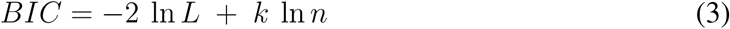

where L is the likelihood, k the number of free parameters, and n the number of single-molecule photon bunches across all experimental conditions. A lower BIC value indicates a statistically better model.^101^ The separation between Gaussian components was assessed using the Ashman’s D, where a score above 2 indicates good separation.^102^ For two Gaussian components with means µ1, µ2 and standard deviations σ1, σ2,

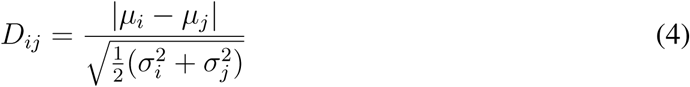

where D*_ij_* represents the distance metric between Gaussian components *i* and *j*. All fitted parameters, likelihood values, BIC scores, and Ashman’s D values are summarized in Supplementary Table 5.

### Statistical information

Statistical analysis was performed using MATLAB. One-way analysis of variance (ANOVA) was performed on different pairs of experimental data and statistical significance was set at P ≤ 0.001. The P-values, degrees of freedom, F-statistics is reported in Supplementary Table 1, 3, 7, 10. The number of data points in the smFRET lifetime distributions is reported in Supplementary Table 2, 4, 8, 11. The number of data points are from two to four biological replicates for each sample.

### Coarse-grained, Explicit-solvent Simulations with the MARTINI Force Field

We conducted a series of analyses based on comprehensive explicit-solvent simulations of the full-length EGFR with the coarse-grained MARTINI force field.^103^ Since the original Martini force field^103^ was parameterized for ordered proteins and will over collapse the disordered C-terminal tail of EGFR, we calibrated the force field by adjusting the protein-water interactions to capture more accurately the size of the C-terminal tail region.^18^ Simulations were performed with the homology-modeled^104^ full EGFR embedded in a lipid bilayer.^105, 106^ 400 µs, including four sets of simulations, were carried out for active/inactive EGFR embedded in the DMPC/POPC-POPS membranes, respectively. Following the typical Martini protocol,^106^ we used a time step of 20 fs, and a temperature of 303 K in all of our simulations. We further used the umbrella sampling technique^107^ to enhance the exploration of the EGFR conformational space, biasing two collective variables: the contact number between the KD domain and the C-terminal tail, and the distance between the JMA domain and the N-terminal part of the CTT domain. Details of the simulation are described in a previous publication.^18^

We analyzed our simulations using WHAM^108, 109^ to reweight the umbrella biases and compute the average values of various metrics introduced in this manuscript. Specifically, we calculated the distance between Residue 721 and Residue 1186 (EGFR C-terminus) of the protein. To quantify the accessibility of the ATP-binding site, we calculated the number of contacts between lipid molecules and the residues forming the ATP-binding pocket (residues 694-703, 719, 766-769, 772-773, 817, 820, and 831).^110^ Close contact between the bilayer and these residues would sterically hinder ATP binding; thus, the contact number serves as a proxy for ATP-site accessibility. The cutoff distance for defining a contact was set to 16 Å, corresponding to the largest molecular radius of the fluorescent ATP analogue (atto647N-ATP, 16.96 Å ^111^). Accordingly, we defined a contact as a pair of coarse-grained atoms, one from the lipid membrane and one from the ATP binding site, within a mutual distance of less than 16 Å.

## Acknowledgments

This work was supported by the NIH MIRA R35 GM148287-02 (to G.S.S.-C.). X.L. and B.Z. acknowledge support by startup funds from the Department of Chemistry at the Massachusetts Institute of Technology. We thank Dr. Adam W. Smith for his valuable feedback on the manuscript.

## Author contributions

S.S. and G.S.S.-C conceived the experiments. S.S. optimized labeled EGFR production in different membrane environments and performed the ensemble fluorescence experiments. X.C. performed the EGFR function-related and characterization experiments. S.S. and R.R performed the single-molecule fluorescence experiments, S.S. analyzed the data. X.L. and B.Z. designed and performed the simulations. S.S. and G.S.S.-C. co-wrote the manuscript. All authors discussed the results and commented on the manuscript.

## Data availability statement

The data supporting the finding of the present study need to be deposited in TBD.

## Supplementary information

**Supplementary Fig. 1.**
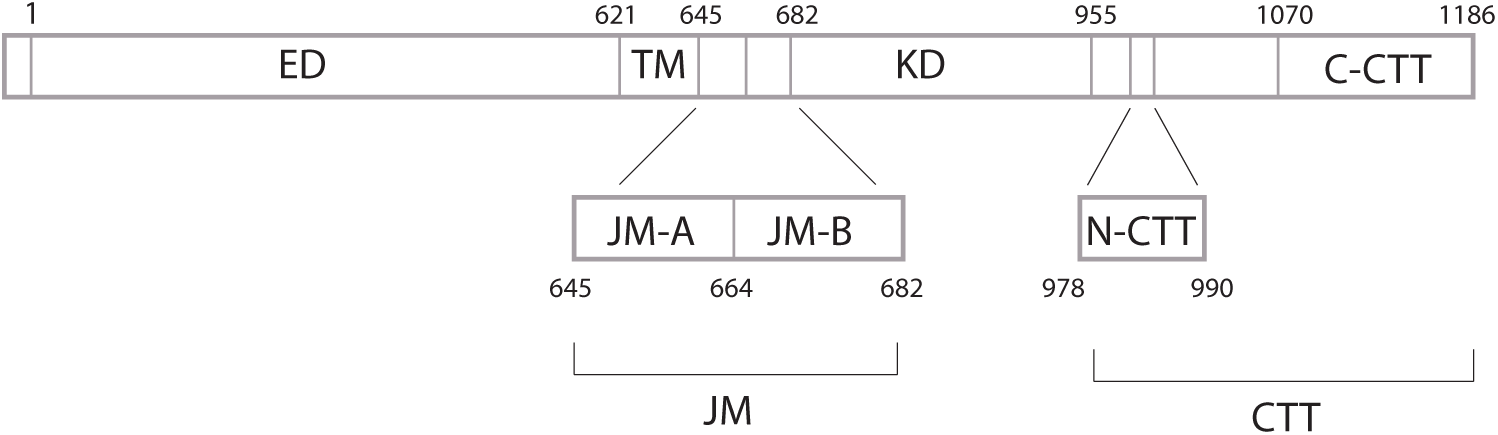
Domains of EGFR. EGFR consists of a 621-amino acid extracellular region (ED), a 24-amino acid transmembrane-spanning domain (TM), and an intracellular region, which is a 37-amino acid juxtamembrane domain (JM), a 273-amino acid kinase domain (KD) and a 231-amino acid C-terminal tail (CTT). The JM is further divided into juxtamembrane-A (JM-A) and juxtamembrane-B (JM-B) domains. Residues 978–990 are defined as N-terminal portion of the CTT (N-CTT) and residues 1070–1186 are defined as the C-terminal portion of the CTT (C-CTT). Residue numbering corresponds to EGFR excluding the signal sequence.

**Supplementary Fig. 2.**
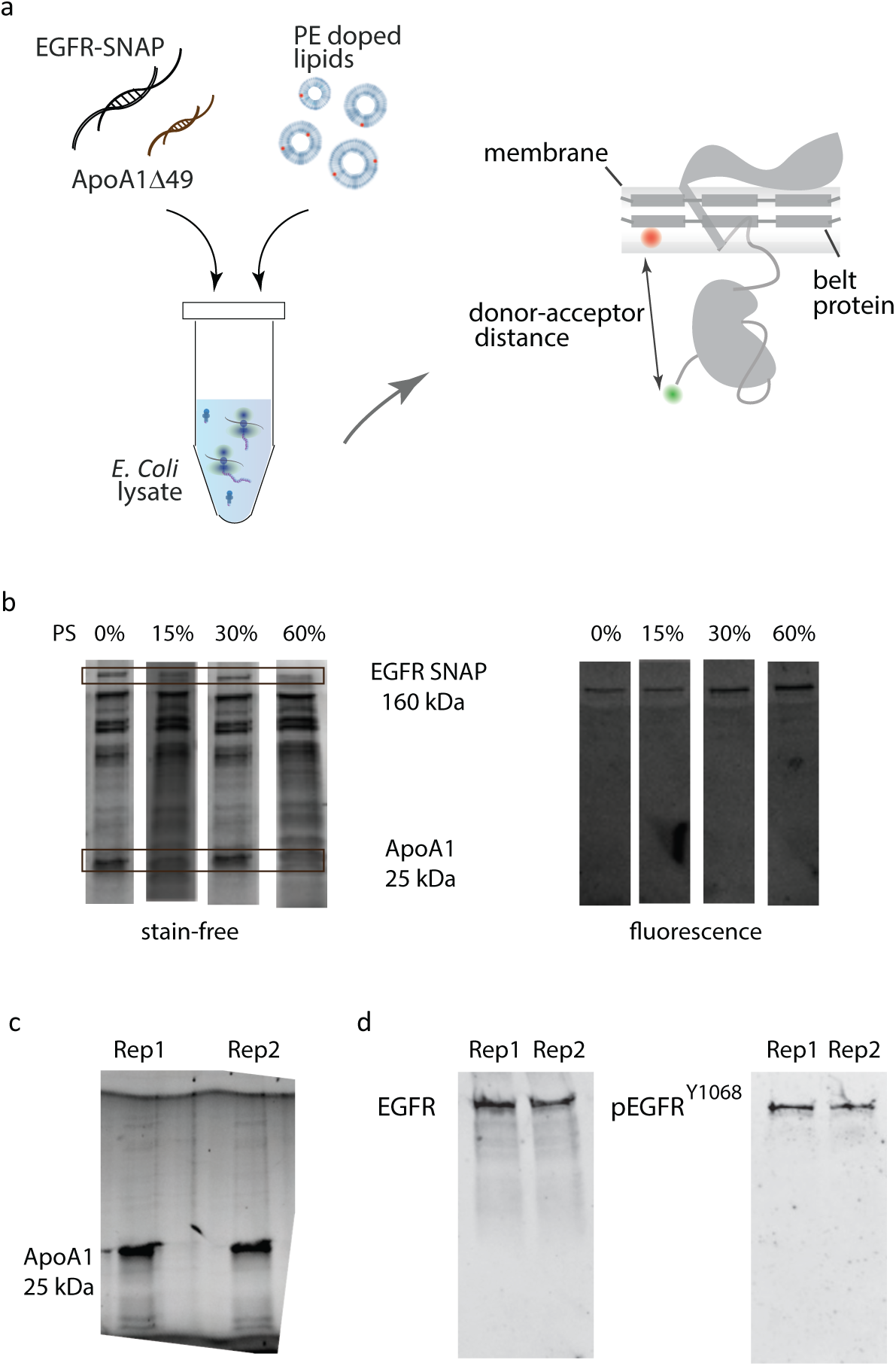
Production and characterization of full-length EGFR in nanodiscs. (a) Cell-free reaction for the production of EGFR and nanodisc belt protein. Codon optimized DNA of the receptor protein (EGFR-SNAP) and the belt protein (ApoA1,6.49) are incubated together with lipid vesicles with or without cholesterol and *E. Coli* lysate at 25*^o^* C. (b) Stain-free (left) and fluorescence (right) gel images of the His-tag purified sample show the presence of ApoA1,6.49 at 25 kDa and full-length EGFR at 160 kDa, which implies successful EGFR production and insertion into nanodiscs. The presence of the EGFR band alone in the fluorescence gel image indicates successful and specific labeling. (c) SDS-PAGE analysis of His-tag purified ApoA1,6.49 (1 µg) produced in cell-free reaction. The two lanes correspond to replicate samples under identical conditions. (d) Western blots were performed on labelled EGFR in nanodiscs. Anti-EGFR Western blots (left) and anti-phosphotyrosine Western blots (right) tested the presence of EGFR and its ability to undergo tyrosine phosphorylation, respectively, consistent with previous experiments on similar preparations.^18,54,55^ The two lanes in each blot correspond to replicate samples under identical conditions.

**Supplementary Fig. 3.**
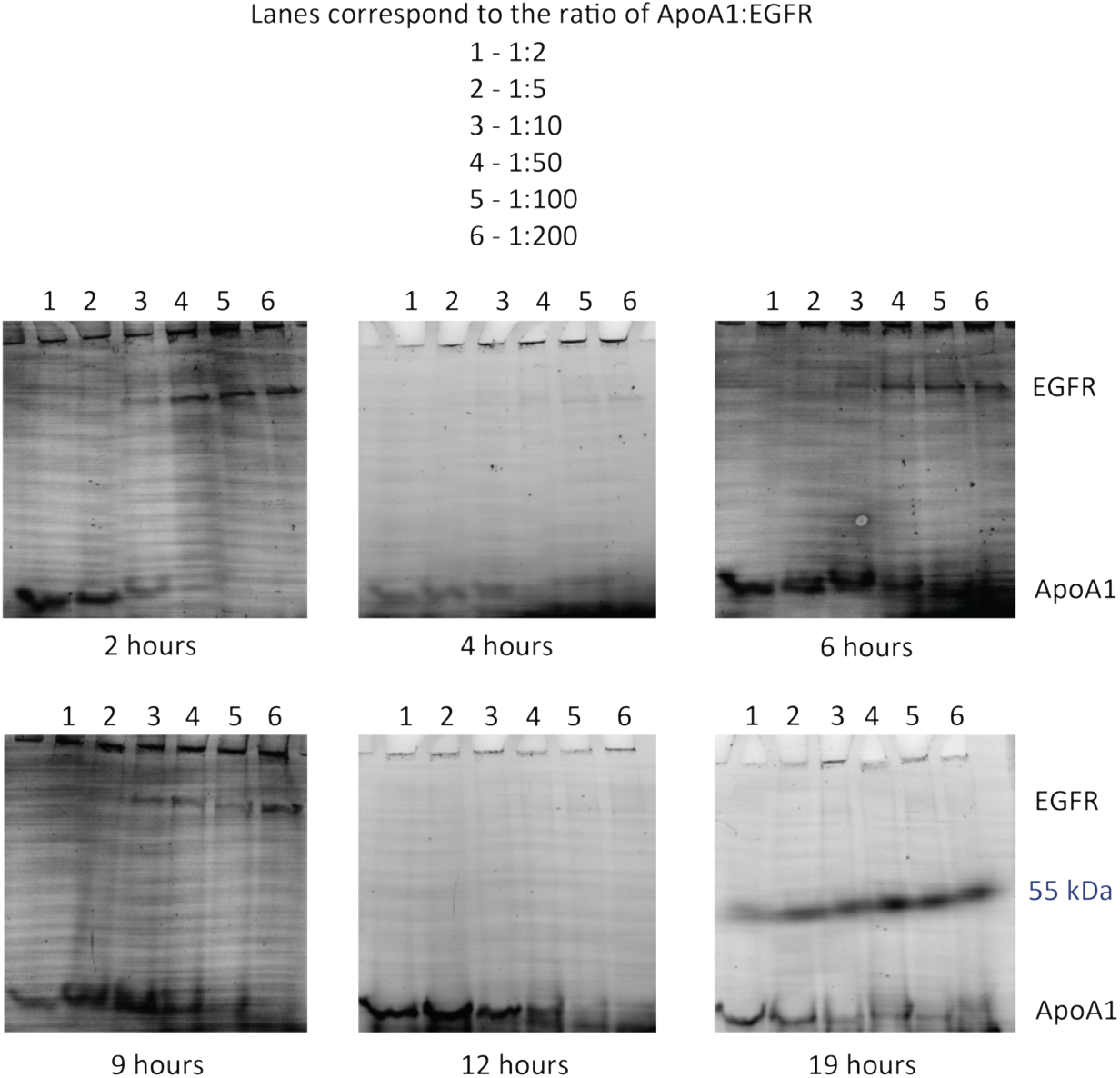
Optimization of ApoA1 and EGFR co-expression in cell-free reactions. Cell-free reactions were performed with different template rations of ApoA1,6.49:EGFR (1:2 to 1:200) and sampled at various time points (2 - 19 hours). Labeled lysines were incorporated during synthesis, ensuring that only proteins expressed in the cell-free reaction were fluorescently labeled. EGFR (160 kDa) and ApoA1,6.49 (25 kDa) expressions were monitored by SDS-PAGE.

**Supplementary Fig. 4.**
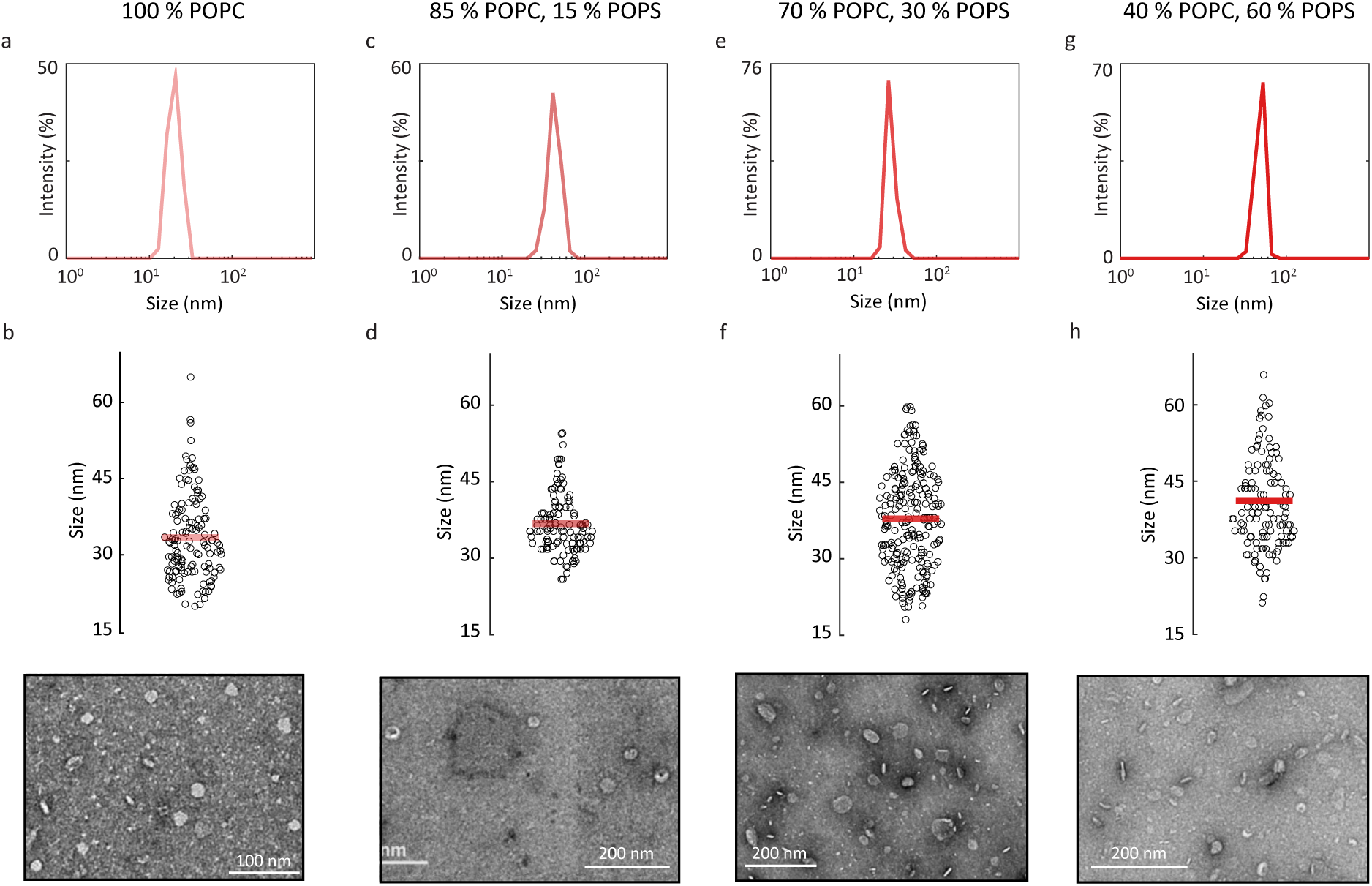
Characterization of EGFR-containing nanodiscs in different anionic membrane environments. (a) Dynamic light scattering (DLS) of EGFR in 100% POPC nanodiscs in PBS buffer indicates ∼38 nm average size. (b) Jitter plot showing the size distribution of EGFR in 100% POPC nanodiscs from negative-stain transmission electron microscopy (TEM). The horizontal line represents the mean size (33.4 *±* 8.5 nm, N = 134). A representative TEM image is shown below. (c) DLS of EGFR in 85% POPC, 15% POPS nanodiscs in PBS buffer indicates ∼40.7 nm average size. (d) Jitter plot showing the size distribution of EGFR in 85% POPC, 15% POPS nanodiscs from negative-stain TEM. The horizontal line represents the mean size (36.8 *±* 6.0 nm, N = 116). A representative TEM image is shown below. (e) DLS of EGFR in 70% POPC, 30% POPS nanodiscs in PBS buffer indicates ∼54.8 nm average size. (f) Jitter plot showing the size distribution of EGFR in 70 % POPC, 30 % POPS nanodiscs from negative-stain TEM. The horizontal line represents the mean size (37.8 *±* 9.7 nm, N = 223). A representative TEM image is shown below. (g) DLS of EGFR in 40 % POPC, 60 % POPS nanodiscs in PBS buffer indicates ∼46.2 nm average size. (h) Jitter plot showing the size distribution of EGFR in 40 % POPC, 60 % POPS nanodiscs from negative-stain TEM. The horizontal line represents the mean size (41.2 *±* 9.7 nm, N = 133). A representative TEM image is shown below.

**Supplementary Fig. 5.**
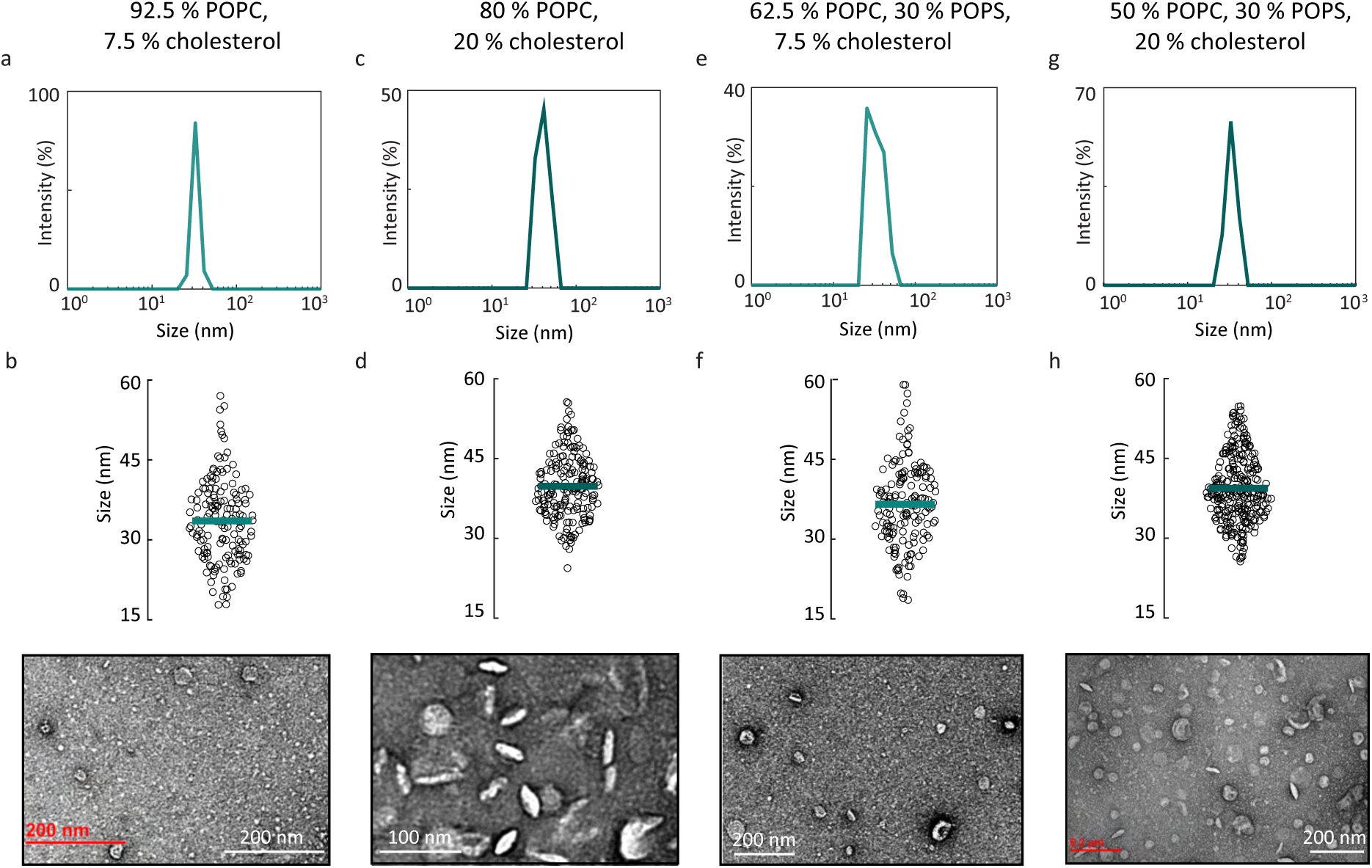
Characterization of EGFR-containing nanodiscs in membrane environments containing cholesterol. (a) Dynamic light scattering (DLS) of EGFR in 92.5% POPC, 7.5% cholesterol containing nanodiscs in PBS buffer indicates indicates 32.8 nm average size. (b) Jitter plot showing the size distribution of EGFR in 92.5% POPC, 7.5% cholesterol nanodiscs from negative-stain transmission electron microscopy (TEM). The horizontal line represents the mean size (33.6 *±* 7.7 nm, N = 154). A representative TEM image is shown below. (c) DLS of EGFR in 80% POPC, 20% cholesterol containing nanodiscs in PBS buffer indicates 39.6 nm average size. (d) Jitter plot showing the size distribution of EGFR in 80% POPC, 20% cholesterol nanodiscs from TEM. The horizontal line represents the mean size (39.8 *±* 5.9 nm, N = 188). A representative TEM image is shown below. (e) DLS of EGFR in EGFR in 62.5% POPC, 30% POPS, 7.5% cholesterol nanodiscs in PBS buffer indicates 33 nm average size. (f) Jitter plot showing the size distribution of EGFR in 62.5% POPC, 30% POPS, 7.5% cholesterol nanodiscs from TEM. The horizontal line represents the mean size (36.5*±* 7.8 nm, N = 159). A representative TEM image is shown below. (g) DLS of EGFR in 50% POPC, 30% POPC, 20% cholesterol nanodiscs in PBS buffer indicates 39.2 nm average size. (h) Jitter plot showing the size distribution of EGFR in 50% POPC, 30% POPC, 20% cholesterol nanodiscs from TEM. The horizontal line represents the mean size (39.2*±* 6.3 nm, N = 281). A representative TEM image is shown below.

**Supplementary Fig. 6.**
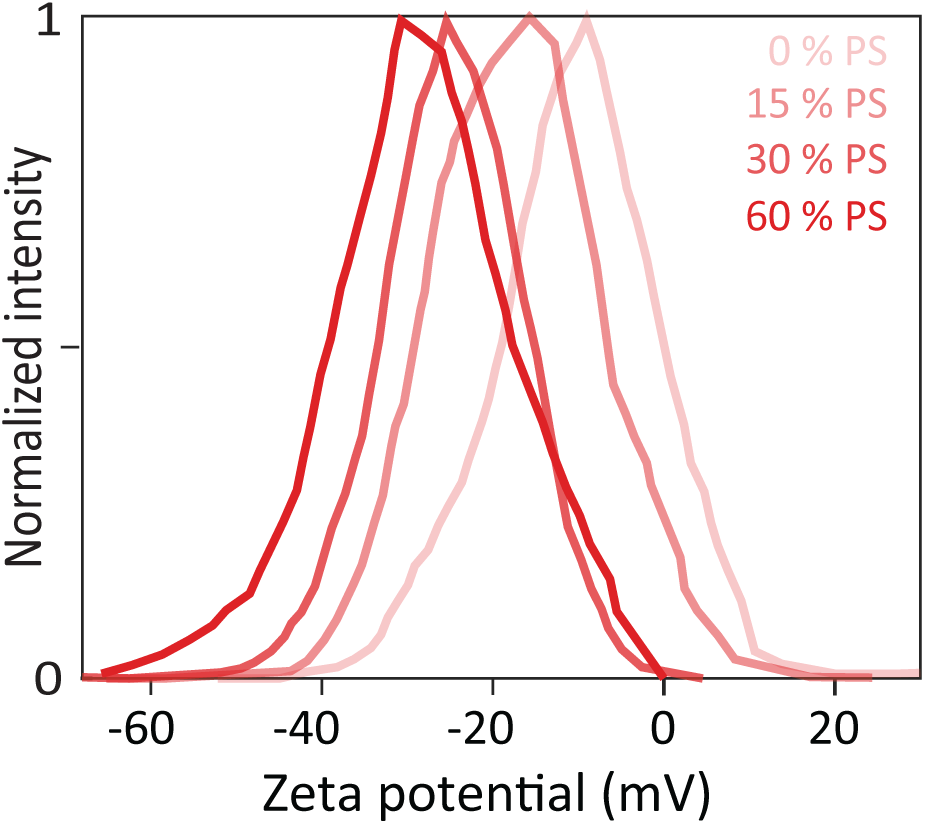
Characterization of anionic content in EGFR-embedded nanodiscs with zeta potential.^56^. Zeta potential distributions for EGFR in nanodiscs containing increasing amounts of anionic lipids (0%, 15%, 30% and 60% POPS). Maximum values from the distributions are shown in Fig. 1b in the main text. Error bars are from three technical replicates.

**Supplementary Fig. 7.**
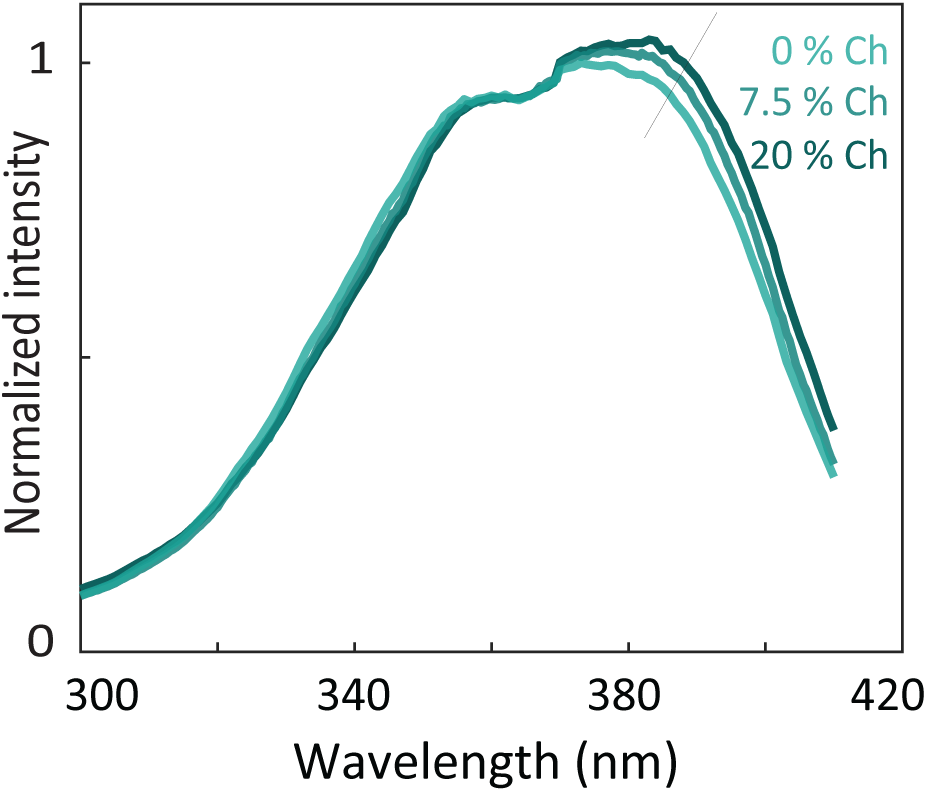
Characterization of cholesterol (Ch) content in EGFR-embedded nanodiscs with Laurdan. Ensemble fluorescence excitation spectra (*λ_em_* = 440 nm) of EGFR embedded Laurdan containing nanodiscs with 0% cholesterol, 7.5% cholesterol and 20% cholesterol. In the Laurdan excitation spectra, increase in the excitation band centered around 390 nm is observed with the addition of increasing amounts of cholesterol to the EGFR embedded nanodisc.^57^

**Supplementary Fig. 8.**
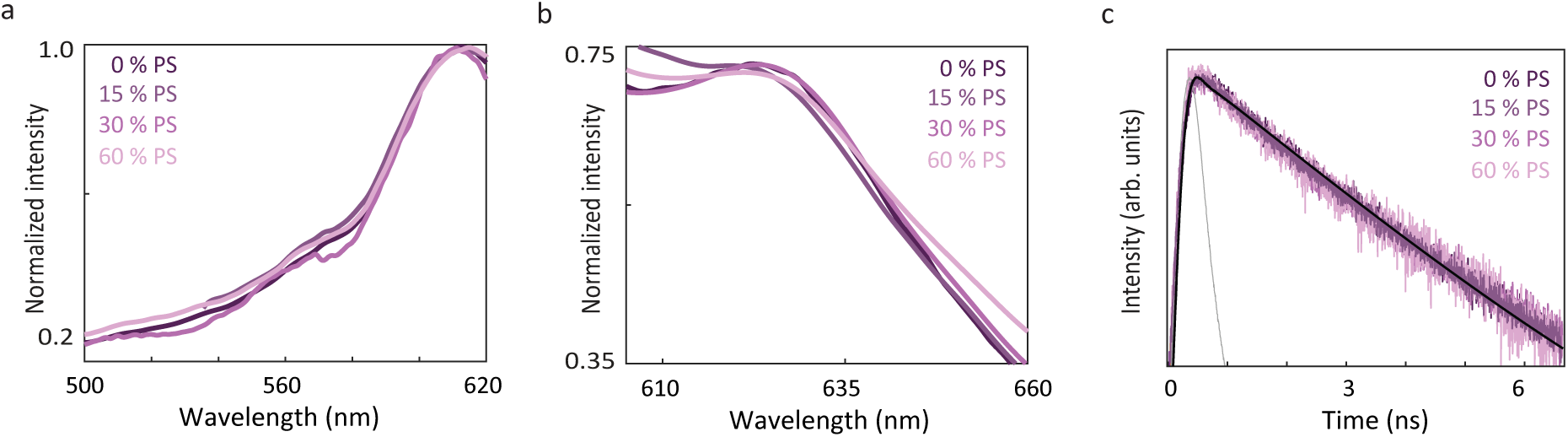
Characterization of ss594 labeled EGFR nanodiscs in different anionic lipid environments. (a) Ensemble fluorescence excitation spectra (*λ_em_* = 660 nm) of ss594 labeled EGFR nanodiscs with 0% POPS, 15% POPS, 30% POPS and 60% POPS lipids. (b) Ensemble fluorescence emission spectra (*λ_exc_* = 565 nm) of ss594 labeled EGFR nanodiscs in 0% POPS, 15% POPS, 30% POPS and 60% POPS lipids. (c) Ensemble time-correlated single photon counting measurements for ss594 labeled EGFR nanodiscs in 0% POPS, 15% POPS, 30% POPS and 60% POPS lipids. The instrument response function (IRF) is shown in gray.

**Supplementary Fig. 9.**
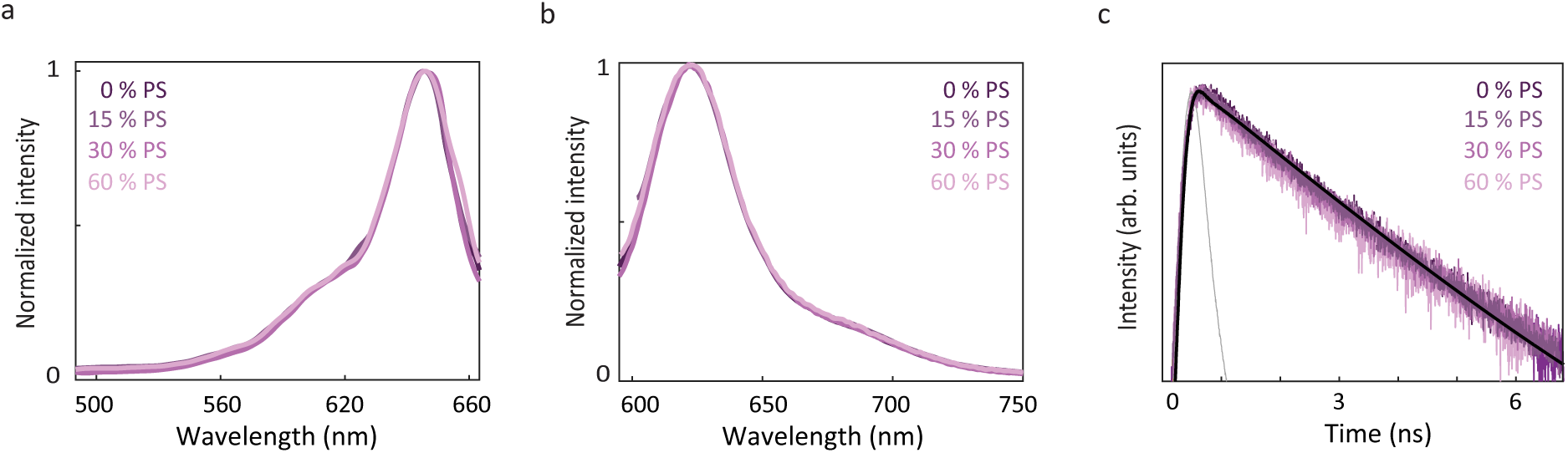
Characterization of cy5 labeled EGFR nanodiscs in different anionic lipid environments. (a) Ensemble fluorescence excitation spectra (*λ_em_* = 700 nm) of cy5 labeled nanodiscs in 0% POPS, 15% POPS, 30% POPS and 60% POPS lipids. (b) Ensemble fluorescence emission spectra (*λ_ex_* = 630 nm) of cy5 labeled nanodiscs in 0% POPS, 15% POPS, 30% POPS and 60% POPS lipids. (c) Ensemble time-correlated single photon counting measurements were performed for cy5 labeled nanodiscs in 0% POPS, 15% POPS, 30% POPS and 60% POPS. The instrument response function (IRF) is shown in gray.

**Supplementary Fig. 10.**
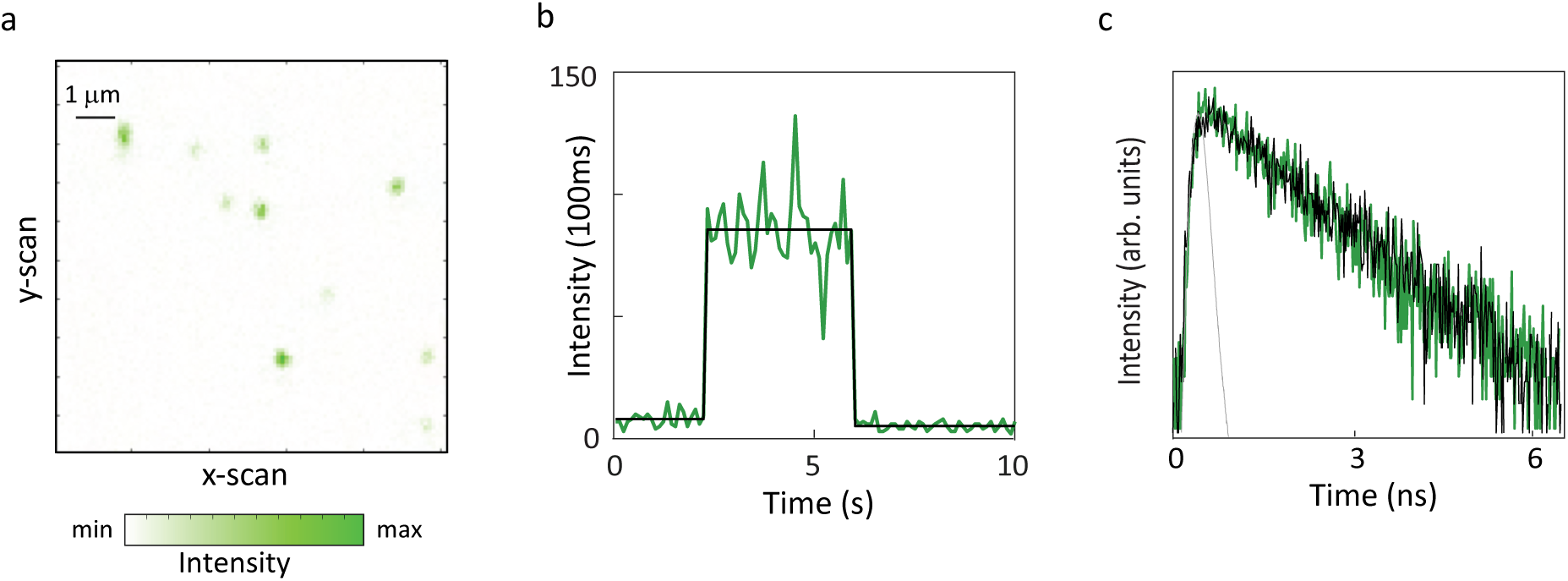
Single ss594 labeled EGFR embedded nanodiscs. (a) Confocal fluorescence image of immobilized constructs of EGFR in nanodiscs labeled with ss594 (*λ_exc_* = 550 nm). (b) Representative intensity time trace from a single construct. The number of detected photons for each 100 ms interval was calculated and used to generate a fluorescence intensity trace (green) with the average intensity for the emissive period overlaid (black). (c) Histogram of the arrival times of detected photons generates the donor lifetime decay profile. Representative decay profiles of EGFR (green) with fit curve (black). The instrument response function (IRF) is shown in gray.

**Supplementary Fig. 11.**
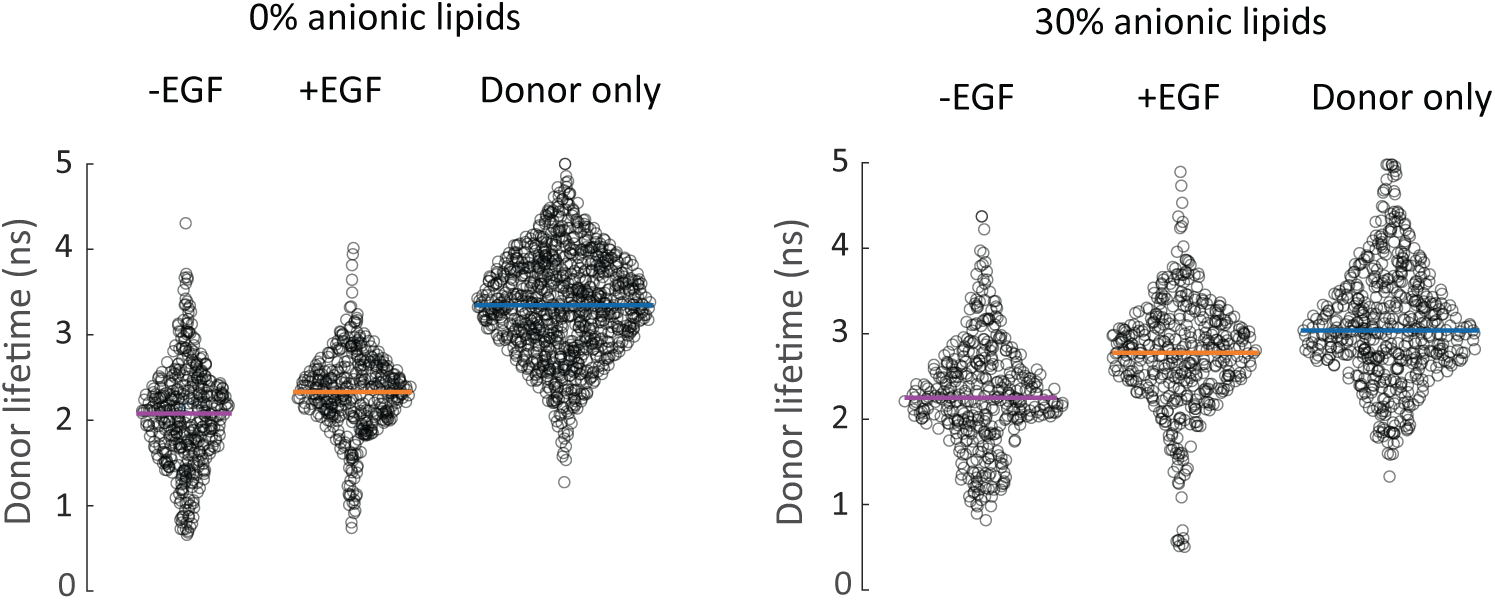
Jitter plots of the donor lifetime distributions from smFRET experiments to measure the distance between the ATP binding site and the C-terminus of EGFR. Distributions of the donor lifetime from the histograms in Fig. 2j, k of main text are represented as jitter plots along with the donor-only samples. The horizontal line indicates the median lifetime. One-way ANOVA was performed to obtain the P-values (Supplementary Table 1). The median value of the donor only distribution in each lipid environment was used as the reference value for calculations of the donor-acceptor distances for all smFRET measurements in that environment.

**Supplementary Fig. 12.**
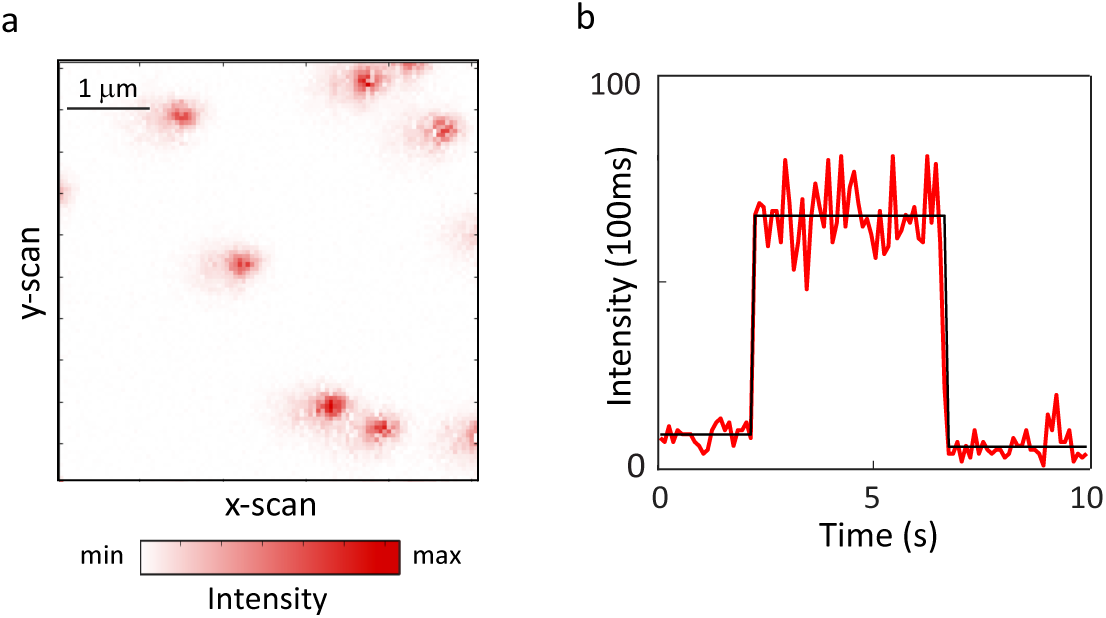
Single Cy5 labeled EGFR embedded nanodiscs. (a) Confocal fluorescence image of immobilized constructs of EGFR in nanodiscs containing a labeled Cy5 lipid (*λ_exc_* = 640 nm). (b) Representative intensity time trace from a single construct. The number of detected photons for each 100 ms interval was calculated and used to generate a fluorescence intensity trace (red) with the average intensity for the emissive period overlaid (black).

**Supplementary Fig. 13.**
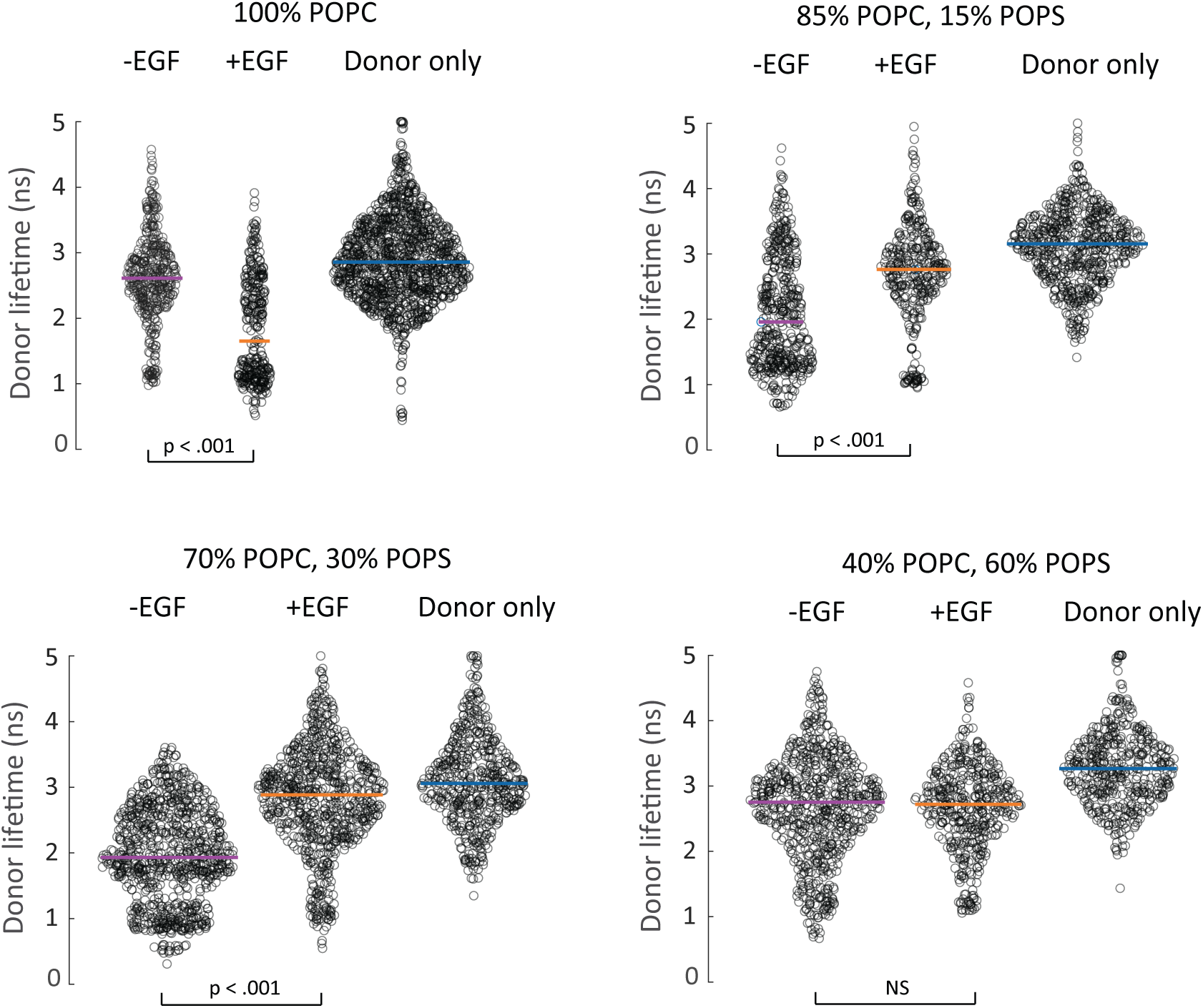
Jitter plots of the donor lifetime distributions from smFRET experiments to measure the distance between the membrane and the C-terminus of EGFR. Distributions of the donor lifetime from the histograms in Fig. 3b of main text are represented as jitter plots along with the donor-only samples. The horizontal line indicates the median lifetime. One-way ANOVA was performed to obtain the P-values (Supplementary Table 3). NS, not significant. The median value of the donor only distribution in each lipid environment was used as the reference value for calculations of the donor-acceptor distances for all smFRET measurements in that environment.

**Supplementary Fig. 14.**
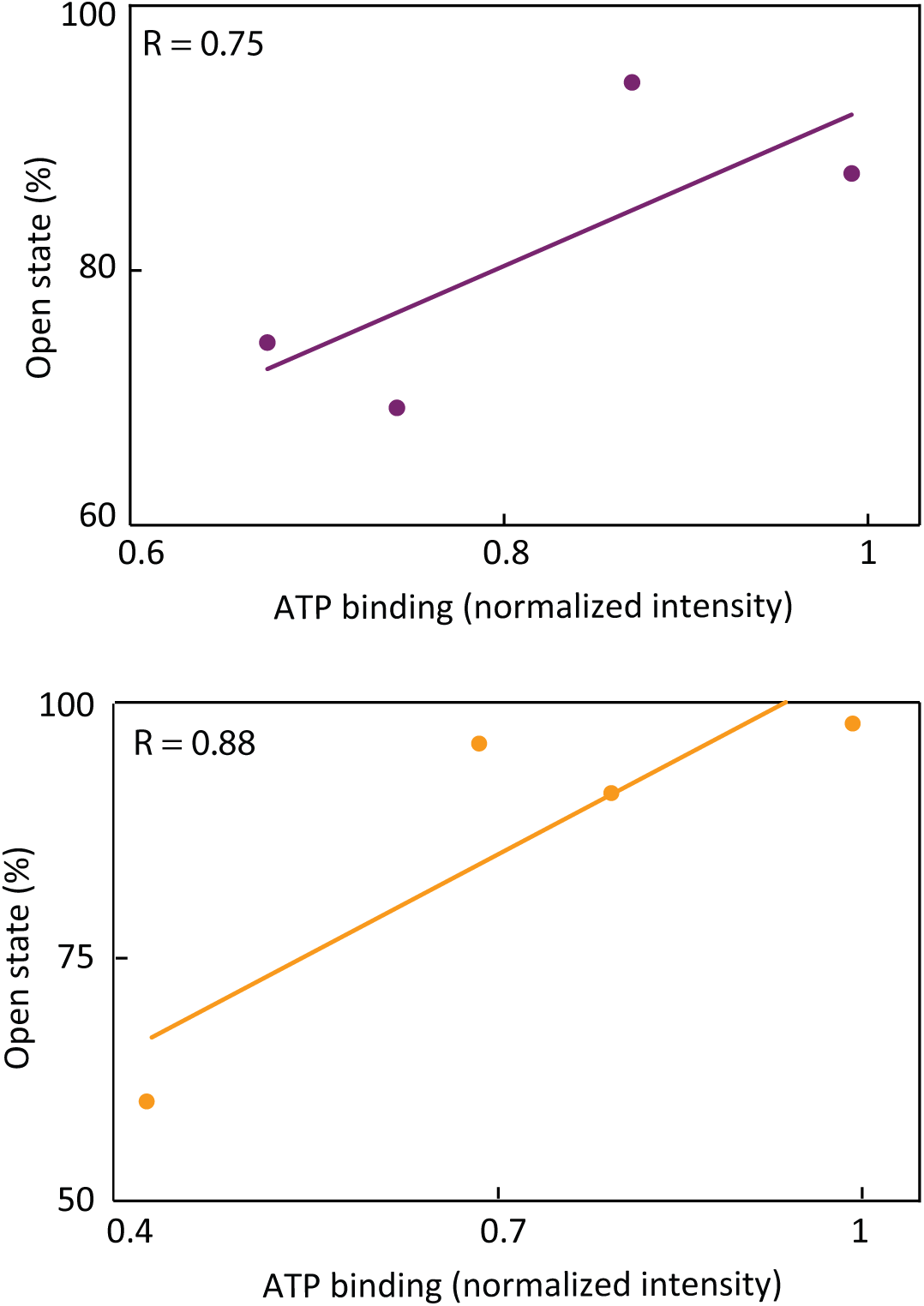
EGFR intracellular domain conformation is correlated with ATP binding. (a) Correlation analysis between distance of EGFR C-terminus from the bilayer and extent of ATP binding in the (top) absence of EGF (Pearson correlation R = 0.75) and (bottom) presence of EGF (Pearson correlation R = 0.88).

**Supplementary Fig. 15.**
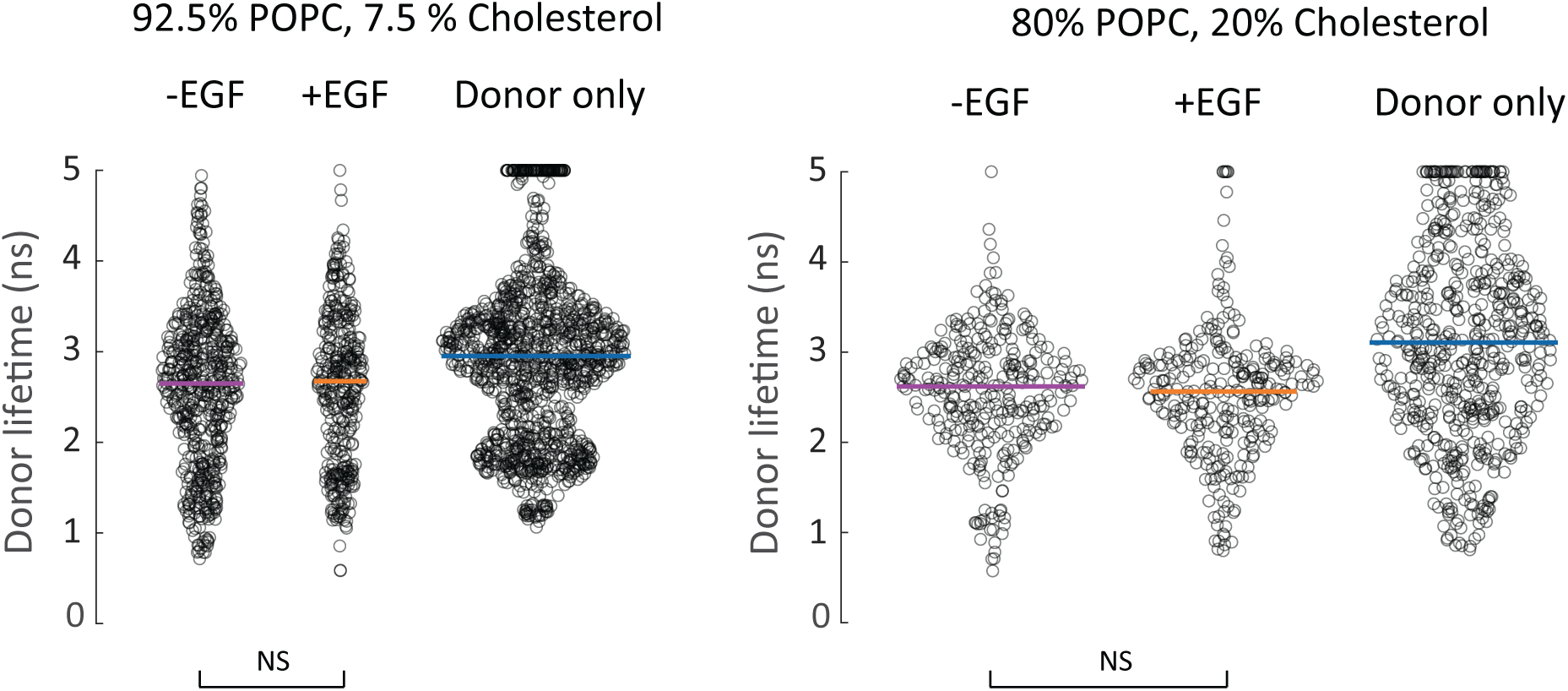
Jitter plots of the donor lifetime distributions from smFRET experiments to measure the distance between the membrane and the C-terminus of EGFR. Distributions of the donor lifetime from the histograms in Fig. 3d of main text are represented as jitter plots along with the donor-only samples. The horizontal line indicates the median lifetime. One-way ANOVA was performed to obtain the P-values (Supplementary Table 7). NS, not significant. The median value of the donor only distribution in each lipid environment was used as the reference value for calculations of the donor-acceptor distances for all smFRET measurements in that environment.

**Supplementary Fig. 16.**
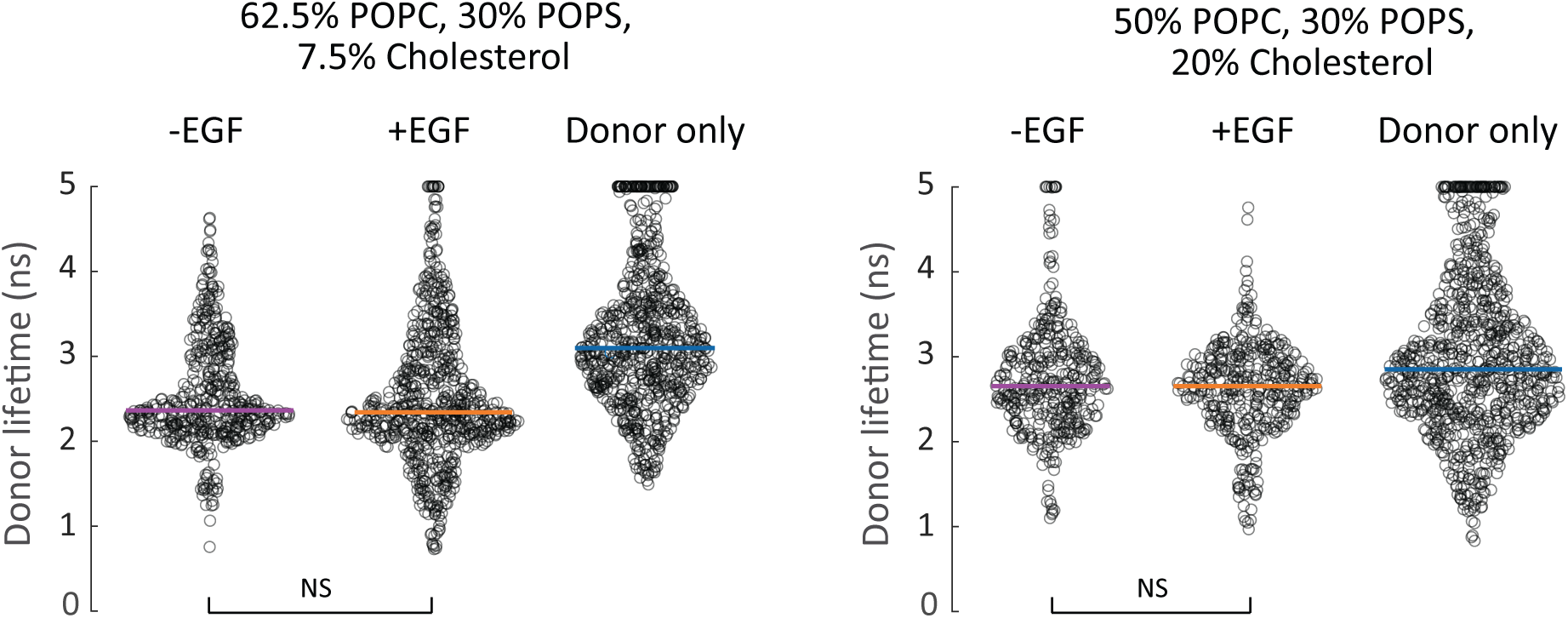
Jitter plots of the donor lifetime distributions from smFRET experiments to measure the distance between the membrane and the C-terminus of EGFR. Distributions of the donor lifetime from the histograms in Fig. 3f of main text are represented as jitter plots along with the donor-only samples. The horizontal line indicates the median lifetime. One-way ANOVA was performed to obtain the P-values (Supplementary Table 10). NS, not significant. The median value of the donor only distribution in each lipid environment was used as the reference value for calculations of the donor-acceptor distances for all smFRET measurements in that environment.

**Supplementary Fig. 17.**
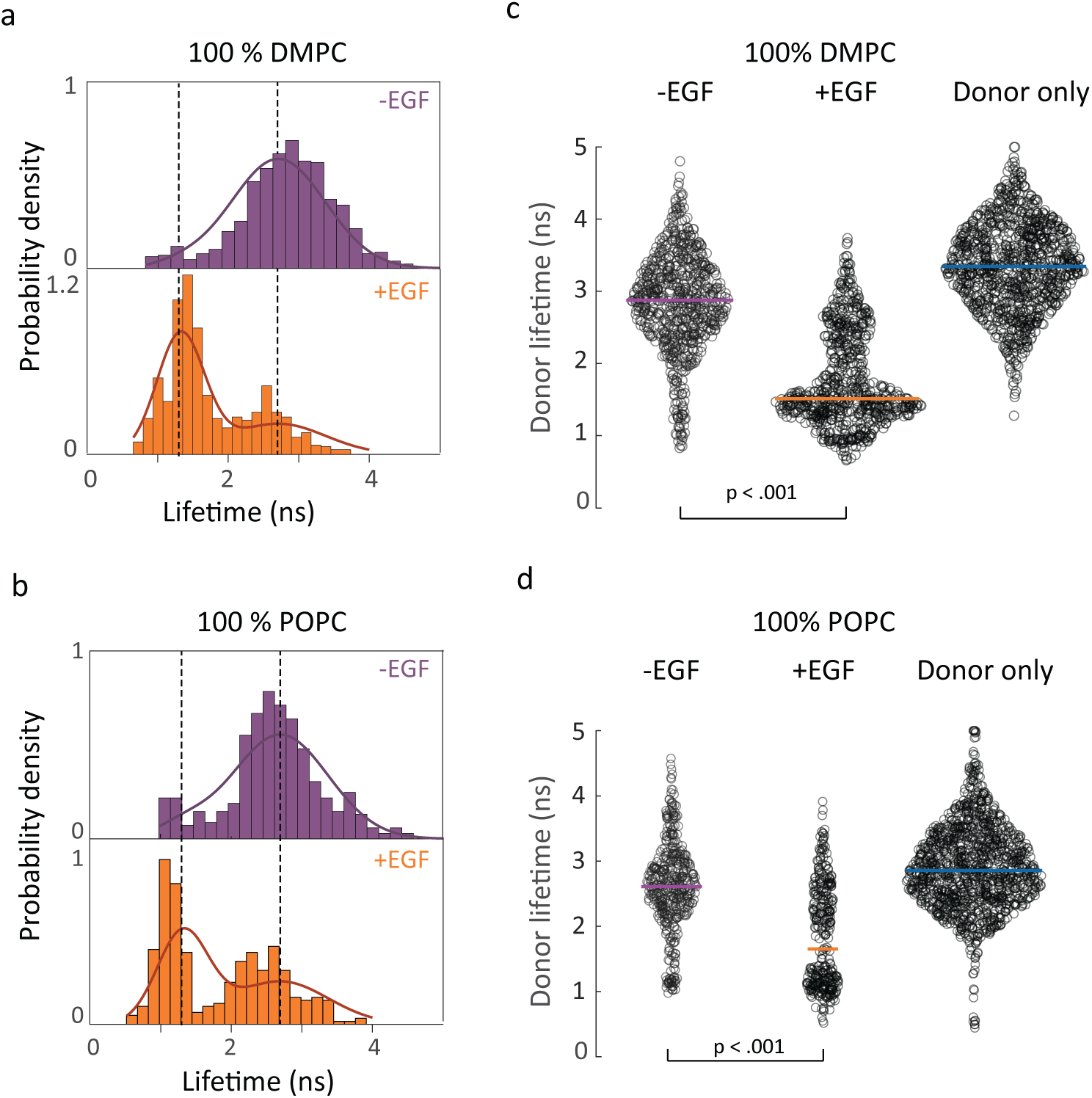
smFRET donor lifetime distributions in DMPC and POPC nanodiscs. smFRET donor fluorescence lifetime distributions in (a) 100% DMPC and (b) 100% POPC without EGF (top); with 1 *µ*M EGF (bottom). Distributions of the donor lifetime from the histograms on the left in 100% DMPC (c) and 100% POPC (d) are represented as jitter plots along with the donor-only samples. The horizontal line indicates the median lifetime. One-way ANOVA was performed to obtain the P-values (Supplementary Table 3). The median value of the donor only distribution in each lipid environment was used as the reference value for calculations of the donor-acceptor distances for all smFRET measurements in that environment.

**Supplementary Table 1.** Statistical analysis of smFRET lifetime distributions. Results from one-way analysis of variance (ANOVA) with P-value, F-statistic and degrees of freedom for all experimental pairs (experiment 1 and experiment 2 in the above table) from the distributions reported in Fig. 2j, k in the main text.

| Sample condition 1 | Sample condition 2 | P-value | F | Degrees of freedom |
| --- | --- | --- | --- | --- |
| 0 % anionic lipids |  |  |  |  |
| EGFR, -EGF | EGFR, +EGF | < 0.001 | 49.4 | 875 |
| 30 % anionic lipids |  |  |  |  |
| EGFR, -EGF | EGFR, +EGF | < 0.001 | 108.8 | 863 |

**Supplementary Table 2.** Sample sizes for smFRET measurements. The number of molecules and number of photon bunches used to construct the lifetime distributions are reported for all smFRET histograms from Fig. 2j, k in the main text.

| Sample conditions | Number of molecules | Number of bunches |
| --- | --- | --- |
| 0 % anionic lipids |  |  |
| EGFR, -EGF | 111 | 448 |
| EGFR, +EGF | 92 | 428 |
| 30 % anionic lipids |  |  |
| EGFR, -EGF | 95 | 422 |
| EGFR, +EGF | 111 | 442 |

**Supplementary Table 3.** Statistical analysis of smFRET lifetime distributions. Results from one-way analysis of variance (ANOVA) with P-value, F-statistic and degrees of freedom for all experimental pairs (experiment 1 and experiment 2 in the above table) from the distributions reported in Fig. 3b in the main text and Supplementary Fig. 17.

| Sample condition 1 | Sample condition 2 | P-value | F | Degrees of freedom |
| --- | --- | --- | --- | --- |
| 0 % POPS, 100 % DMPC;<br>-EGF | 0 % POPS, 100 % DMPC;<br>+EGF | < 0.001 | 911.3 | 1398 |
| 0 % POPS, 100 % POPC;<br>-EGF | 0 % POPS, 100 % POPC;<br>+EGF | < 0.001 | 204.48 | 757 |
| 15 % POPS, 85 % POPC;<br>-EGF | 15 % POPS, 85 % POPC;<br>+EGF | < 0.001 | 104.87 | 850 |
| 30 % POPS, 70 % POPC;<br>-EGF | 30 % POPS, 70 % POPC;<br>+EGF | < 0.001 | 540.13 | 1686 |
| 60 % POPS, 40 % POPC;<br>-EGF | 60 % POPS, 40 % POPC;<br>+EGF | 0.7512 | 0.1 | 1193 |

**Supplementary Table 4.** Sample sizes for smFRET measurements. The number of molecules and number of photon bunches used to construct the lifetime distributions are reported for all smFRET histograms from Fig. 3b in the main text and Supplementary Fig. 17.

| Sample conditions | Number of molecules | Number of bunches |
| --- | --- | --- |
| 0 % POPS, 100 % DMPC; -EGF | 51 | 702 |
| 0 % POPS, 100 % DMPC; +EGF | 53 | 697 |
| 0 % POPS, 100 % DMPC; donor-only | 77 | 992 |
| 0 % POPS, 100 % POPC; -EGF | 74 | 422 |
| 0 % POPS, 100 % POPC; +EGF | 75 | 336 |
| 0 % POPS, 100 % POPC; donor-only | 122 | 1254 |
| 15 % POPS, 85 % POPC; -EGF | 87 | 490 |
| 15 % POPS, 85 % POPC; +EGF | 76 | 361 |
| 15 % POPS, 85 % POPC; donor-only | 107 | 665 |
| 30 % POPS, 70 % POPC; -EGF | 155 | 891 |
| 30 % POPS, 70 % POPC; +EGF | 118 | 796 |
| 30 % POPS, 70 % POPC; donor-only | 59 | 551 |
| 60 % POPS, 40 % POPC; -EGF | 150 | 723 |
| 60 % POPS, 40 % POPC; +EGF | 71 | 471 |
| 60 % POPS, 40 % POPC; donor-only | 113 | 488 |

**Supplementary Table 5.**
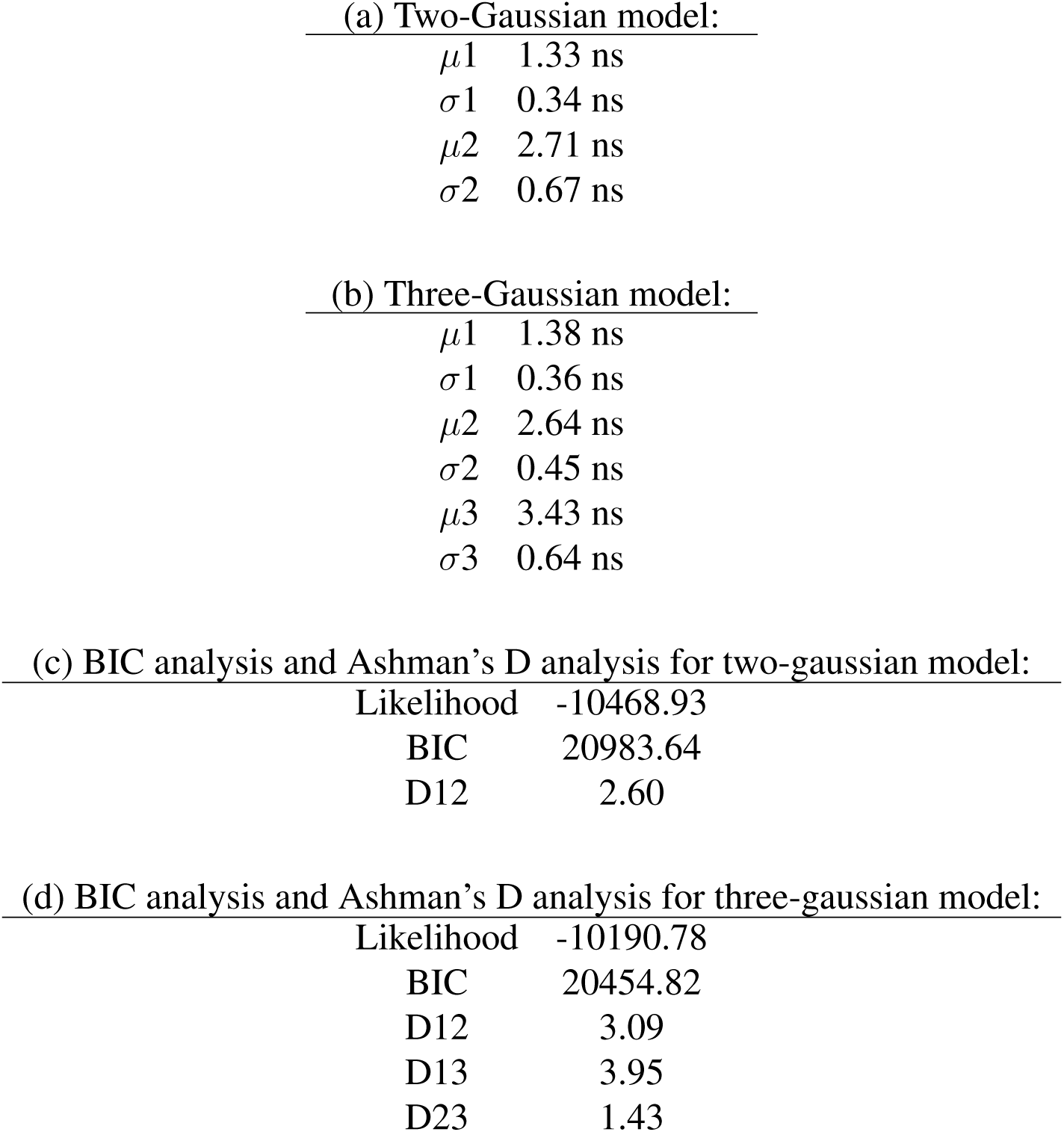
Supplementary Table 1. Model selection and statistical analysis of global fits. Comparison of two- and three-Gaussian models used to globally fit the lifetime distributions across all experimental conditions. Parameters (*µ*, *o*) correspond to the mean and standard deviation of each Gaussian component, respectively. Likelihood, Bayesian Information Criterion (BIC), and Ashman’s D values are reported for assessing model quality and separation of components. The number of free parameters is 5 for the two-Gaussian model and 8 for the three-Gaussian model. The number of single-molecule photon bunches = 9,483 across all 18 conditions (lipid composition and *±*EGF).

|  |  |
| --- | --- |
| $\mu 1$ | 1.33 ns |
| $\sigma 1$ | 0.34 ns |
| $\mu 2$ | 2.71 ns |
| $\sigma 2$ | 0.67 ns |
| (b) Three-Gaussian model: |  |
| $\mu 1$ | 1.38 ns |
| $\sigma 1$ | 0.36 ns |
| $\mu 2$ | 2.64 ns |
| $\sigma 2$ | 0.45 ns |
| $\mu 3$ | 3.43 ns |
| $\sigma 3$ | 0.64 ns |
| (c) BIC analysis and Ashman's D analysis for two-gaussian model: |  |
| Likelihood | -10468.93 |
| BIC | 20983.64 |
| D12 | 2.60 |

**Supplementary Table 5. Supplementary Table 1.**
|  |  |
| --- | --- |
| Likelihood | -10190.78 |
| BIC | 20454.82 |
| D12 | 3.09 |
| D13 | 3.95 |
| D23 | 1.43 |

**Supplementary Table 6.**
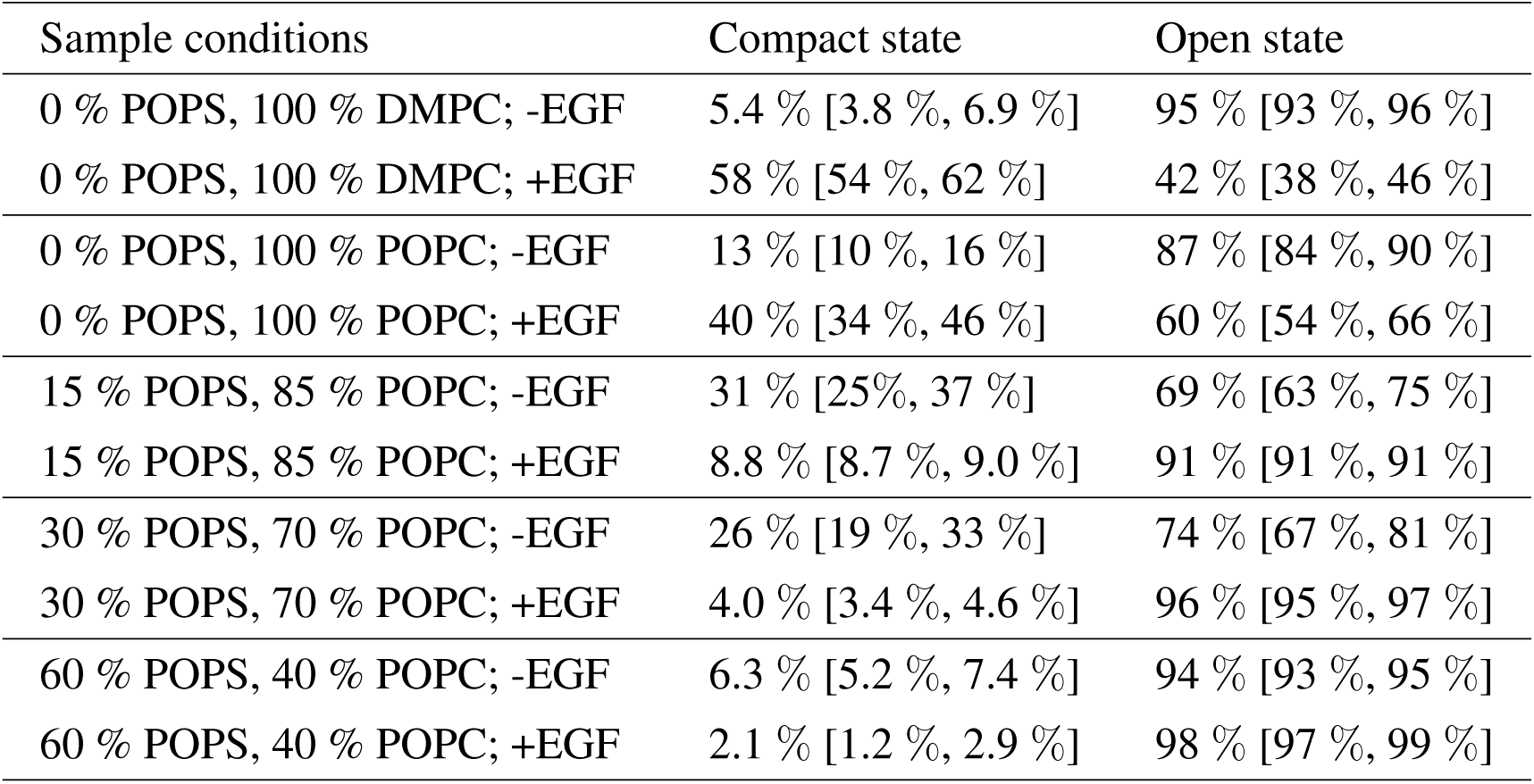
Amplitude of compact and open state of the EGFR intracellular domain from smFRET lifetime distributions in main text Fig. 3b. The numbers in parenthesis indicates the error bar indicated in Fig. 3g.

| Sample conditions | Compact state | Open state |
| --- | --- | --- |
| 0 % POPS, 100 % DMPC; -EGF | 5.4 % [3.8 %, 6.9 %] | 95 % [93 %, 96 %] |
| 0 % POPS, 100 % DMPC; +EGF | 58 % [54 %, 62 %] | 42 % [38 %, 46 %] |
| 0 % POPS, 100 % POPC; -EGF | 13 % [10 %, 16 %] | 87 % [84 %, 90 %] |
| 0 % POPS, 100 % POPC; +EGF | 40 % [34 %, 46 %] | 60 % [54 %, 66 %] |
| 15 % POPS, 85 % POPC; -EGF | 31 % [25%, 37 %] | 69 % [63 %, 75 %] |
| 15 % POPS, 85 % POPC; +EGF | 8.8 % [8.7 %, 9.0 %] | 91 % [91 %, 91 %] |
| 30 % POPS, 70 % POPC; -EGF | 26 % [19 %, 33 %] | 74 % [67 %, 81 %] |
| 30 % POPS, 70 % POPC; +EGF | 4.0 % [3.4 %, 4.6 %] | 96 % [95 %, 97 %] |
| 60 % POPS, 40 % POPC; -EGF | 6.3 % [5.2 %, 7.4 %] | 94 % [93 %, 95 %] |
| 60 % POPS, 40 % POPC; +EGF | 2.1 % [1.2 %, 2.9 %] | 98 % [97 %, 99 %] |

**Supplementary Table 7.** Statistical analysis of smFRET lifetime distributions. Results from one-way analysis of variance (ANOVA) with P-value, F-statistic and degrees of freedom for all experimental pairs (experiment 1 and experiment 2 in the above table) from the distributions reported in Fig. 3d in the main text.

| Sample condition 1 | Sample condition 2 | P-value | F | Degrees of freedom |
| --- | --- | --- | --- | --- |
| 92.5 % POPC,<br>7.5 % Cholesterol;<br>-EGF | 92.5 % POPC,<br>7.5 % Cholesterol;<br>+EGF | 0.91 | 0.01 | 975 |
| 80 % POPC,<br>20 % Cholesterol;<br>-EGF | 80 % POPC,<br>20 % Cholesterol;<br>+EGF | 0.8245 | 0.05 | 600 |

**Supplementary Table 8.**
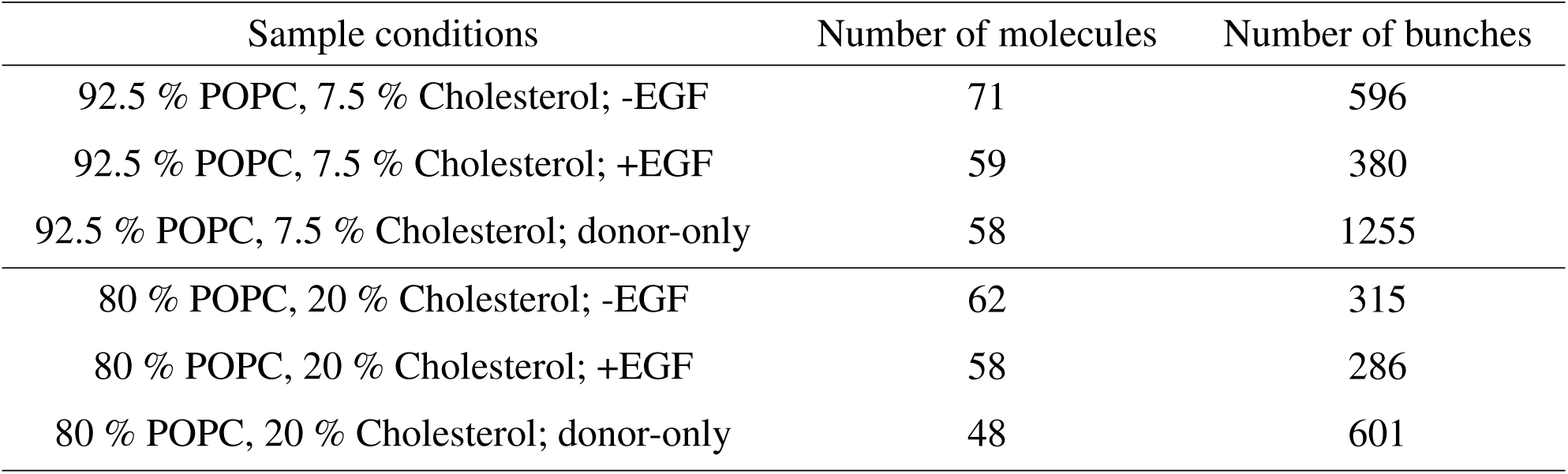
Sample sizes for smFRET measurements. The number of molecules and number of photon bunches used to construct the lifetime distributions are reported for all smFRET histograms from Fig. 3d in the main text.

**Supplementary Table 9.**
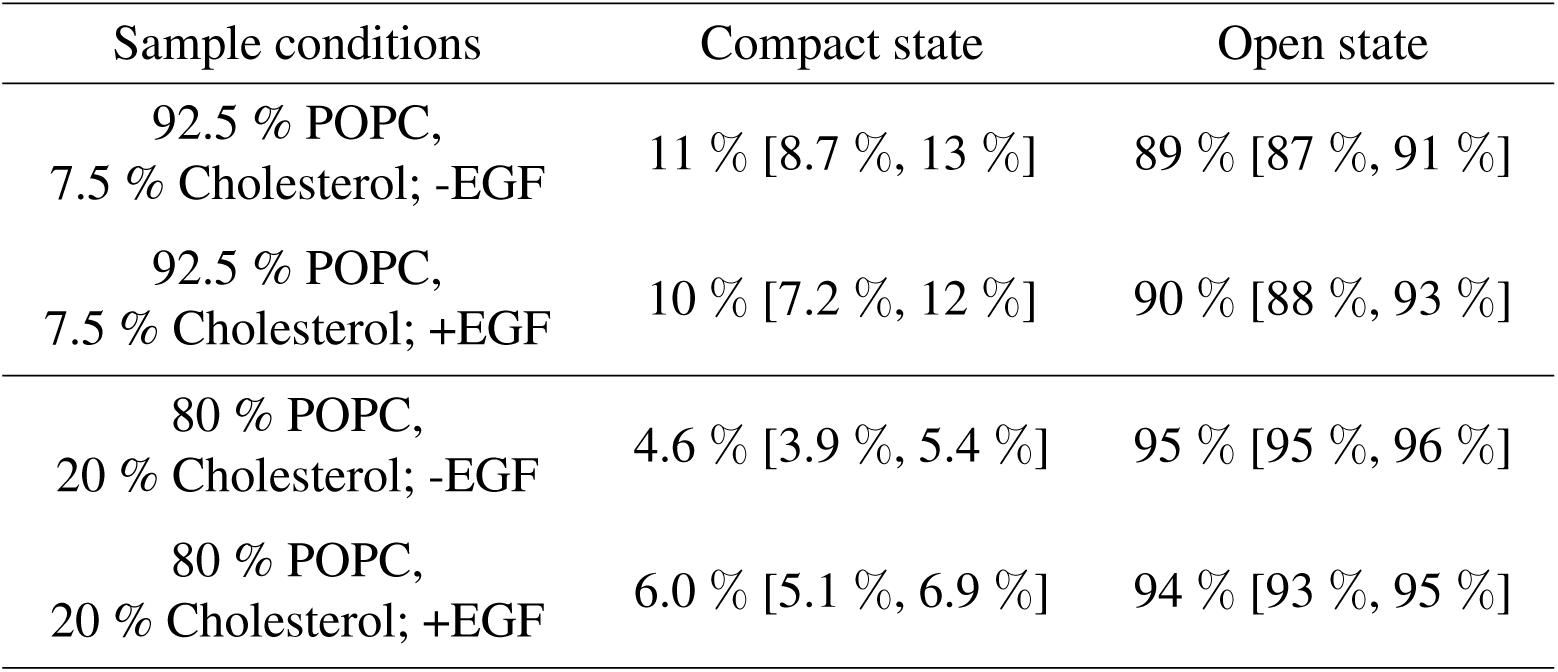
Amplitude of compact and open state of the EGFR intracellular domain from smFRET lifetime distributions in main text Fig. 3d. The numbers in parenthesis indicates the error bar indicated in Fig. 3g.

| Sample conditions | Compact state | Open state |
| --- | --- | --- |
| 92.5 % POPC,<br>7.5 % Cholesterol; -EGF | 11 % [8.7 %, 13 %] | 89 % [87 %, 91 %] |
| 92.5 % POPC,<br>7.5 % Cholesterol; +EGF | 10 % [7.2 %, 12 %] | 90 % [88 %, 93 %] |
| 80 % POPC,<br>20 % Cholesterol; -EGF | 4.6 % [3.9 %, 5.4 %] | 95 % [95 %, 96 %] |
| 80 % POPC,<br>20 % Cholesterol; +EGF | 6.0 % [5.1 %, 6.9 %] | 94 % [93 %, 95 %] |

**Supplementary Table 10.** Statistical analysis of smFRET lifetime distributions. Results from one-way analysis of variance (ANOVA) with P-value, F-statistic and degrees of freedom for all experimental pairs (experiment 1 and experiment 2 in the above table) from the distributions reported in Fig. 3f in the main text.

| Sample condition 1 | Sample condition 2 | P-value | F | Degrees of freedom |
| --- | --- | --- | --- | --- |
| 62.5 % POPC,<br>30 % POPS,<br>7.5 % Cholesterol,<br>-EGF | 62.5 % POPC,<br>30 % POPS,<br>7.5 % Cholesterol,<br>+EGF | 0.61 | 0.27 | 1203 |
| 50 % POPC,<br>30 % POPS,<br>20 % Cholesterol,<br>-EGF | 50 % POPC,<br>30 % POPS,<br>20 % Cholesterol,<br>+EGF | 0.002 | 9.88 | 794 |

**Supplementary Table 11.**
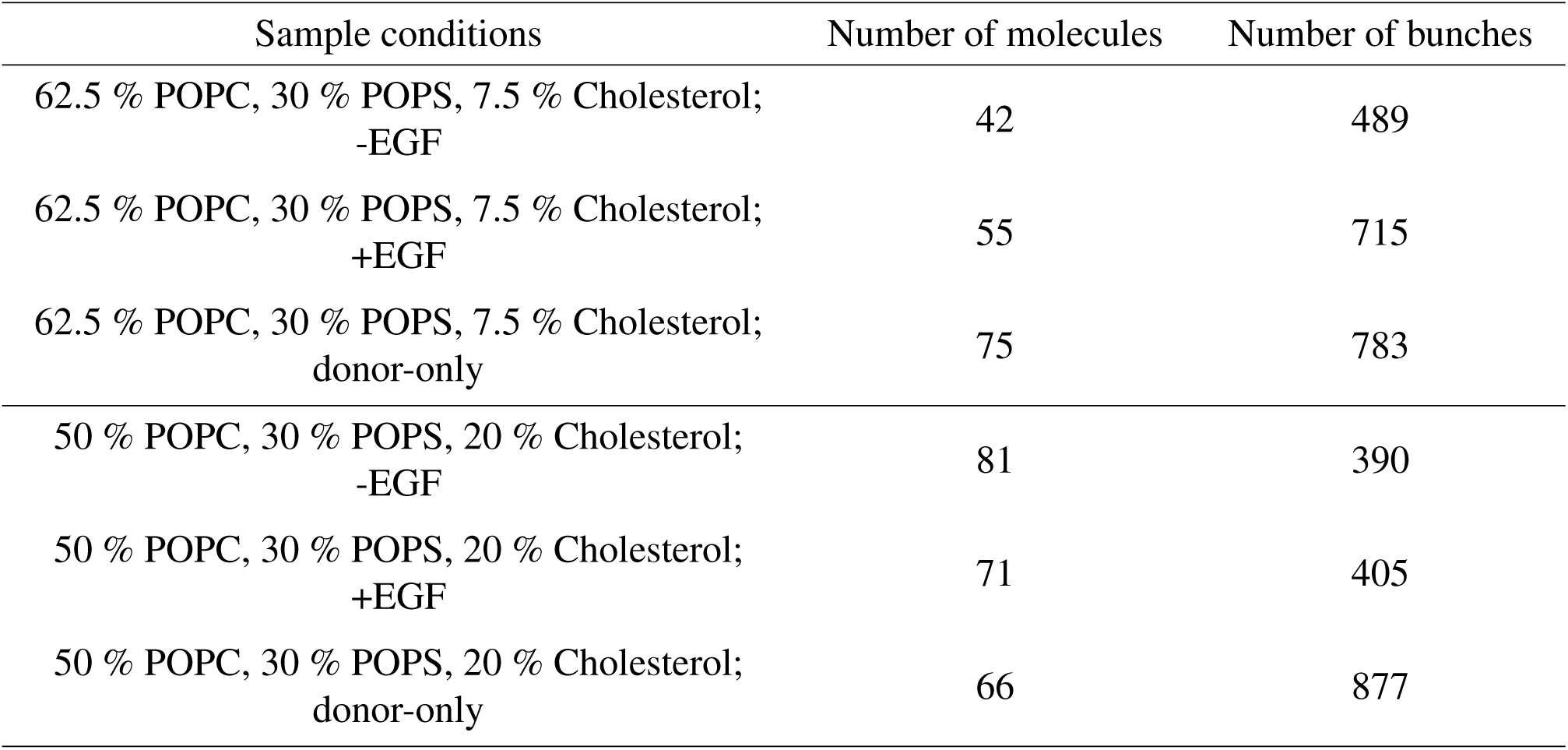
Sample sizes for smFRET measurements. The number of molecules and number of photon bunches used to construct the lifetime distributions are reported for all smFRET histograms from Fig. 3f in the main text.

**Supplementary Table 12.** Amplitude of compact and open state of the EGFR intracellular domain from smFRET lifetime distributions in main text Fig. 3f. The numbers in parenthesis indicates the error bar indicated in Fig. 3g.

| Sample conditions | Compact state | Open state |
| --- | --- | --- |
| 62.5 % POPC, 30 % POPS,<br>7.5 % Cholesterol, -EGF | 0.0 % [0.0 %, 0.0 %] | 100 % [100 %, 100 %] |
| 62.5 % POPC, 30 % POPS,<br>7.5 % Cholesterol, +EGF | 5.9 % [4.8 %, 7.0 %] | 94 % [93 %, 95 %] |
| 50 % POPC, 30 % POPS,<br>20 % Cholesterol, -EGF | 0 % [0 %, 0 %] | 100 % [100 %, 100 %] |
| 50 % POPC, 30 % POPS,<br>20 % Cholesterol, +EGF | 0.9 % [0.8 %, 1.0 %] | 99 % [99 %, 99 %] |

**Supplementary Table 13.** List of antibodies used to show phosphorylation of EGFR nanodiscs.

| Antibody name | Target | Host | Company | Clone | Dilution |
| --- | --- | --- | --- | --- | --- |
| Anti-EGFR Antibody<br>(A-10) | EGFR<br>C-terminal | MouseIgG<sub>2a</sub> | Santa Cruz<br>Biotechnology | Monoclonal | 1:200 |
| Human Phospho-EGFR<br>Y1068 Antibody | EGFR<br>phosphorylated<br>at Y1068 | MouseIgG<sub>2a</sub> | R&D Systems | Monoclonal | 1:200 |
| Goat anti-Mouse<br>IgG (H+L) Highly Cross-Adsorbed Antibody<br>Alexa Fluor 790 | Mouse | GoatIgG | Thermo Fisher | Polyclonal | 1:10000 |

